# T cells promote distinct transcriptional programs of cutaneous inflammatory disease in keratinocytes and dermal fibroblasts

**DOI:** 10.1101/2024.07.31.606077

**Authors:** Hannah A. DeBerg, Mitch L. Fahning, Suraj R. Varkhande, James D. Schlenker, William P. Schmitt, Aayush Gupta, Archana Singh, Iris K. Gratz, Jeffrey S. Carlin, Daniel J. Campbell, Peter A. Morawski

**Affiliations:** Center for Systems Immunology, Benaroya Research Institute, Seattle, WA, USA; Center for Fundamental Immunology, Benaroya Research Institute, Seattle, WA, USA; Department of Biosciences and Medical Biology, University of Salzburg, Salzburg, Austria; Plastic and Reconstructive Surgery, Virginia Mason Medical Center, Seattle, WA, USA; Department of Dermatology, Leprology, and Venereology, Dr. D. Y. Patil Medical College, Hospital and Research Centre, Pune, India; Systems Biology Lab, CSIR – Institute of Genomics and Integrative Biology, New Delhi, India; Academy of Scientific and Innovative Research (AcSIR), Gaziabad, India; EB House Austria, Department of Dermatology, University Hospital of the Paracelsus Medical University, Salzburg, Austria; Center for Tumor Biology and Immunology, University of Salzburg, Salzburg, Austria; Center for Translational Immunology, Benaroya Research Institute, Seattle, WA, USA; Division of Rheumatology, Virginia Mason Medical Center, Seattle, WA, USA; Department of Immunology, University of Washington School of Medicine, Seattle, WA, USA

**Author notes:** Correspondence: 1201 9th Ave, Seattle, WA 98101, USA.

## Abstract

T cells and structural cells coordinate appropriate inflammatory responses and restoration of barrier integrity following insult. Dysfunctional T cells precipitate skin pathology occurring alongside altered structural cell frequencies and transcriptional states, but to what extent different T cells promote disease-associated changes remains unclear. We show that functionally diverse circulating and skin-resident CD4^+^CLA^+^ T cell populations promote distinct transcriptional outcomes in human keratinocytes and fibroblasts associated with inflamed or healthy tissue. We identify T_h_17 cell-induced genes in keratinocytes that are enriched in psoriasis patient skin and normalized by anti-IL-17 therapy. We also describe a CD103^+^ skin-resident T cell-induced transcriptional module enriched in healthy controls that is diminished during psoriasis and scleroderma and show that CD103^+^ T cell frequencies are altered during disease. Interrogating clinical data using immune-dependent transcriptional signatures defines the T cell subsets and genes distinguishing inflamed from healthy skin and allows investigation of heterogeneous patient responses to biologic therapy.

**GRAPHICAL ABSTRACT:** 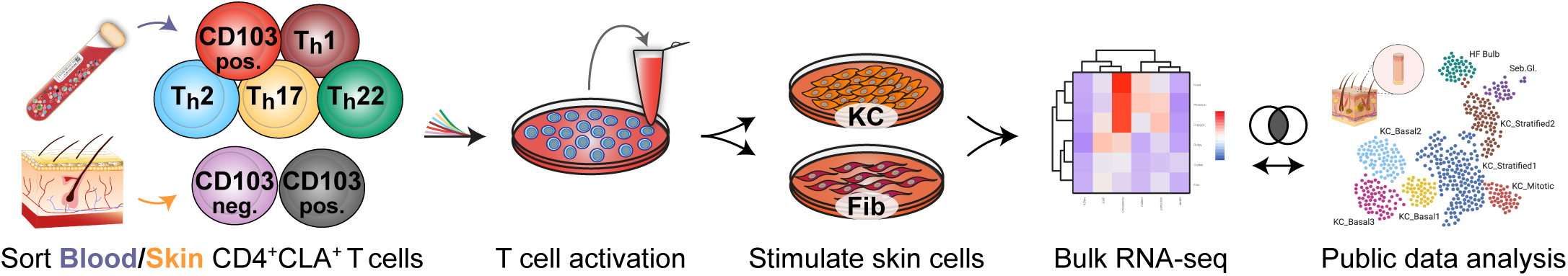

## INTRODUCTION

The skin is an immunologically active barrier tissue specialized to deal with a litany of stresses. It contains many immune and structural cell populations whose frequencies, differentiation, and function are altered during inflammatory skin disease. In the epidermis, numerous distinct keratinocyte (KC) and stem cell states have been described by single cell RNA-seq in healthy skin that are altered during psoriasis (Ps) ^1^. Transcriptional diversity in the dermis is similarly evident and supported by single cell analyses detailing the gene programs of structural cells such as fibroblasts, pericytes, and endothelial cells ^2–5^. One study described the distinct dermal fibroblast (Fib) of healthy subjects and monitored these in individuals with systemic sclerosis (scleroderma; SSc). A progressive loss of LGR5-expressing Fibs (Fib-LGR5), a dominant gene signature in healthy skin, corresponded with SSc severity ^3^. Despite the practical advances made in defining the broad transcriptional diversity of skin structural cells, a conceptual framework on how the distinct gene states of skin are regulated during health and disease is lacking.

Functionally specialized skin T cells cooperate with cells of the epidermis and dermis to respond to chemical, biological, and physical stresses ^6^. Skin T cells are important in the context of responses to pathogens, allergens, and tumors, in barrier maintenance and wound healing, and they drive disease pathogenesis in many autoimmune and inflammatory diseases. Healthy adult human skin has an average surface area of 1.8m^2^ and contains numerically more T cells than any other non-lymphoid tissue: an estimated 2 x 10^10^ T cells or ∼6% of the total T cell mass in the body ^7^. Most skin T cells express cutaneous lymphocyte antigen (CLA), an inducible carbohydrate modification of P-selectin glycoprotein ligand-1 (PSGL-1) and other cell surface glycoproteins ^8,9^. CLA-expressing T cells in the blood are a skin-tropic population expressing markers of tissue homing – CCR4, CCR6, CCR8, CCR10 – and make up 5-15% of the circulating CD4^+^ T cell pool. As with conventional circulating CD4^+^ T cells, CLA^+^ skin-tropic T cells in the blood can be subdivided into distinct helper subsets (T_h_) using standard chemokine receptor identification strategies ^10^. In addition to traditional T_h_ subsets, we previously identified CD4^+^CLA^+^ CD103^+^ T cells, a distinct population of skin resident-memory T cells (T_RM_) that have the capacity to exit the tissue and form a stable fraction of migratory T_RM_ in the circulation of healthy individuals ^11^. We showed that CD4^+^CLA^+^ CD103^neg^ and CD103^+^ skin-resident T cells from healthy donors are transcriptionally, functionally, and clonally distinct populations. Thus, CD4^+^CLA^+^ T cells of the blood and skin include many functionally specialized populations, but how they exert their influence on structural cells of the epidermis and dermis to coordinate skin biology remains incompletely understood.

The contribution of T cells and their cytokines is essential to the development of inflammatory skin disease. Accumulation of T cells in lesional tissue and T cell cytokines in patient serum often serve as indicators of active disease, while GWAS have identified polymorphisms in cytokines and cytokine receptors that increase the risk of developing inflammatory skin disease ^12–14^. Attempts to target cytokine signaling therapeutically in the skin date back four decades to the use of recombinant IFN in cutaneous T cell lymphoma patients ^15^. More recently, IL-17 and IL-23 neutralizing therapies have had immense success at reversing skin pathology during Ps ^16,17^. Many candidate therapeutics aimed at altering the abundance or activity of T cell subsets and cytokines are now being used or tested in patients with inflammatory and autoimmune diseases including Ps, atopic dermatitis (AD, eczema), alopecia areata, hidradenitis suppurativa, and SSc ^18,19^. While clinical interventions targeting T cell activity during inflammatory skin disease are efficacious, it remains unclear why certain individuals are non-responders to treatment and how functionally specialized cutaneous T cell subsets instruct the structural cell gene programs that distinguish diseased skin from healthy tissue.

In this study, we show that subsets of circulating and skin-resident CD4^+^CLA^+^ T cells promote an array of discrete transcriptional outcomes in primary human epithelial KCs and dermal Fibs including the induction of gene programs involved in the inflammatory response, proliferation, barrier integrity, and wound repair. We use this atlas of T cell-dependent effects to investigate gene expression changes occurring during inflammatory skin disease and following intervention in patients receiving anti-cytokine therapy. While some T cell populations potentiate inflammatory responses by KCs and Fibs, others contribute to the maintenance of healthy skin-associated transcriptional programs. These findings inform how T cells contribute to the diverse gene states of skin structural cells, forming the basis for a method to monitor immune-dependent differences between inflamed and healthy tissue and deconvolute patient responses to therapeutic intervention.

## RESULTS

### Functional characterization of circulating and skin-resident human T cell populations

To assess T cell functional capacity, we isolated 7 distinct blood or skin derived CD4^+^CLA^+^ T_h_ subsets (*Blood*: T_h_1, T_h_2, T_h_17, T_h_22, CD103^+^; *Skin*: CD103^neg^, CD103^+^) from healthy controls as described previously ^10,11,20^ (**Figure 1a, 1b**). Sorted T cell populations were stimulated through the TCR and co-stimulatory receptors for 48 hours and the production of 13 common T cell cytokines was measured in culture supernatants (**Figure 1c**, **S1**). As expected, T_h_ cell signature cytokines were enriched: T_h_1 (IFNγ), T_h_2 (IL-4, IL-5, IL-13), T_h_17 (IL-17A, IL-17F), T_h_22 (IL-22), CD103^+^ (IL-13, IL-22). Relative to the blood-derived T cells, those isolated from healthy skin produced lower levels of most measured cytokines (**Figure 1c**). Blood- and skin-derived T cells did both produce comparably high levels of IL-9 (**Figure S1**), a transient signature cytokine of recently activated CLA^+^ T cells ^9^. Thus, CD4^+^CLA^+^ T cell populations of both blood and skin produce an array of signature cytokines associated with host defense, tissue repair, and the development of inflammatory skin disease ^21^.

**Figure 1.**
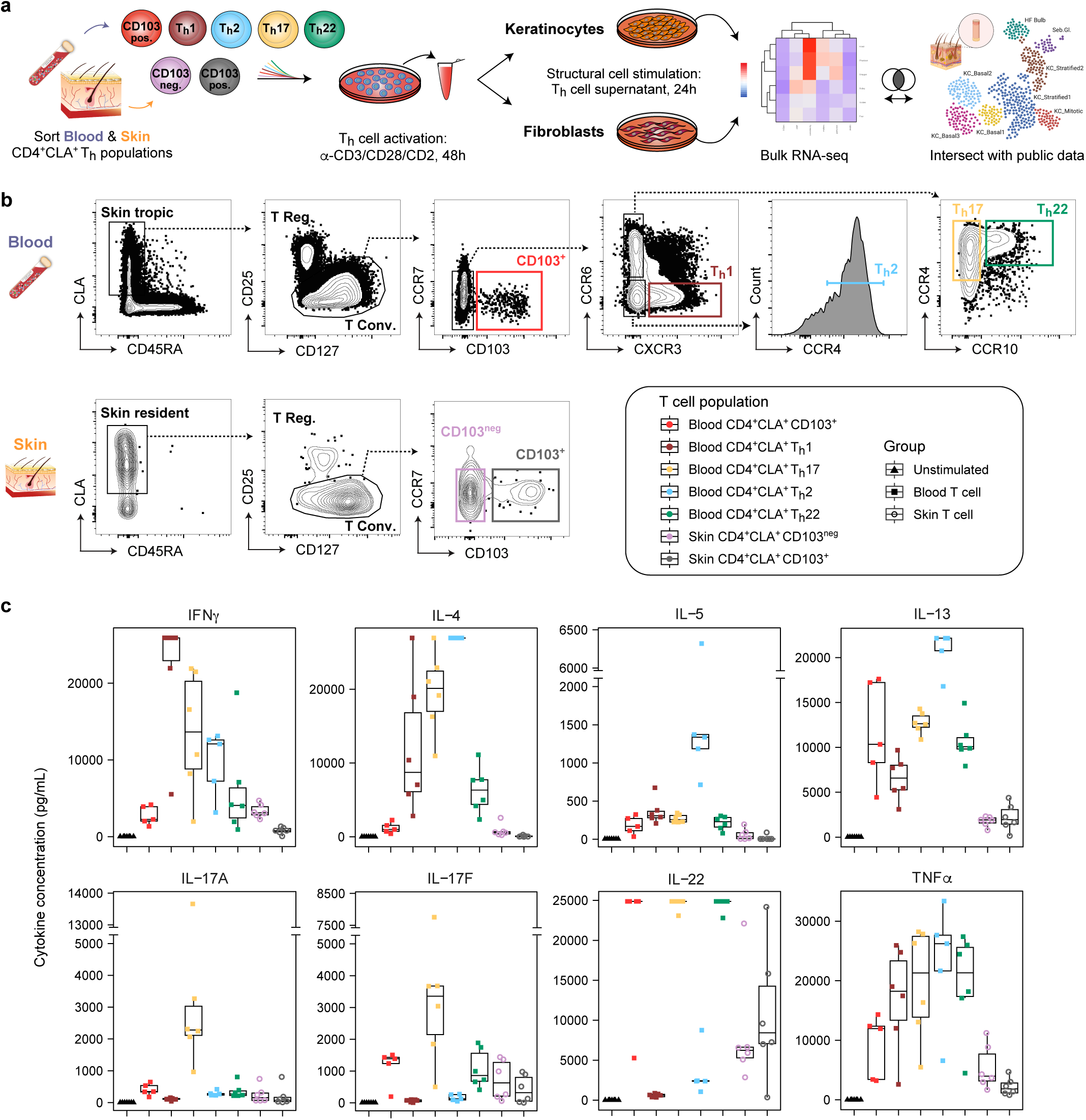
Isolation and functional characterization of circulating and skin-resident human CD4^+^CLA^+^ T cell populations. (**a**) Experimental design and study schematic: Blood and skin CD4^+^CLA^+^ Th cell isolation, activation, and subsequent stimulation of keratinocytes (KC) and fibroblasts (Fib), analysis by bulk RNA-seq, and comparison to public gene expression data from human skin. (**b**) CD4^+^CLA^+^ Th cell sorting strategy for indicated blood and skin populations. T Conv, conventional T cells; T Reg, regulatory T cells. (**c**) Quantitative cytokine bead array measurement of stimulated T cell supernatants for the 8 indicated analytes (n=5-6 donors per population). Full statistical analysis of cytokine production is provided in **Table S1**.

### CD4^+^CLA^+^ T cells induce distinct transcriptional states in epithelial keratinocytes and dermal fibroblasts

Assessing the effects of individual cytokines on the skin has uncovered specific aspects of tissue biology, but this approach fails to recreate the complexity of an inflammatory response. To determine how each CD4^+^CLA^+^ T_h_ cell subset impacts the gene expression profile of skin structural cells, we cultured T cell-stimulated supernatants from 5-7 healthy donors per sorted tissue group (blood, skin) together with healthy donor primary KCs or Fibs for 24 hours. The transcriptional response was then assessed by bulk RNA-seq (**Figure 1a**). Unsupervised principal component (PC) analysis revealed that T cell tissue of origin and T_h_ cell population are the major sources of variance in treated KCs (**Figure 2a, 2b**). Each CD4^+^CLA^+^ T cell population promoted a distinct transcriptional state in the KCs. The top differentially expressed (DE) genes in KCs across all conditions clustered according to both the T cell subset and its tissue of origin (**Figure 2c**). Patterns in T cell-induced gene expression corresponded with the concentrations of cytokines measured in the supernatant of those T cells (IFNγ, IL-4, IL-5, IL-13, IL-17A, IL17F, IL-22, TNFα). A subset of inflammatory response genes known to be induced by T_h_1 (i.e. IDO1, CXCL11), T_h_2 (i.e. CCL26, IL13RA2), and T_h_17 cells (i.e. CSF3, IL23A) also showed expression patterns directly related to the detected amount of IFNγ, IL-4/IL-5/IL-13, or IL-17A/F, respectively (**Figure 2d**). By comparison, skin derived CD103^+^ T cells and to a lesser extent the migratory CD103^+^ fraction from the blood drove the expression of genes associated with cell cycle and proliferation in KCs (i.e. MKI67, CDK1).

**Figure 2.**
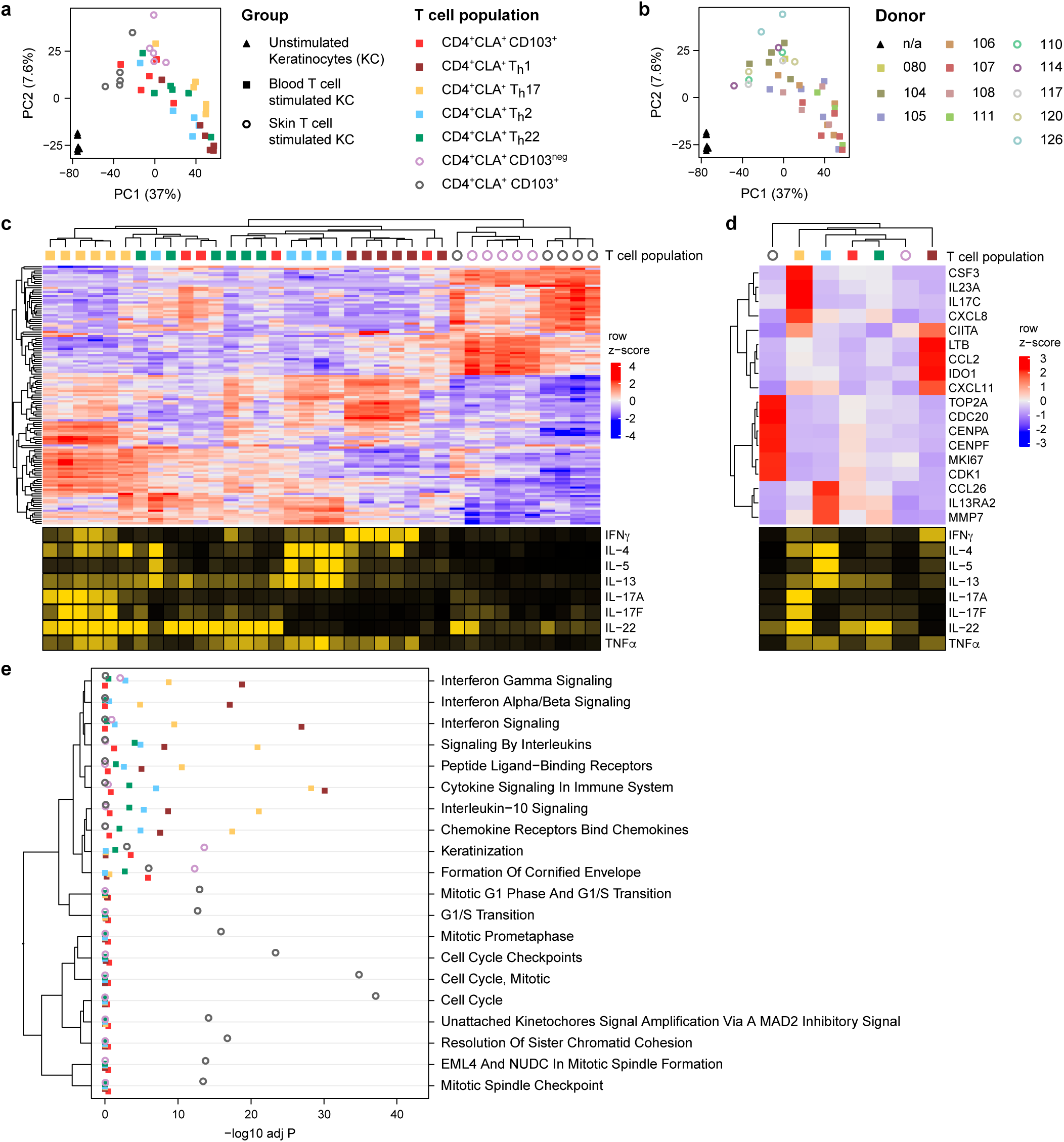
CD4^+^CLA^+^ T cells induce distinct transcriptional states in epithelial keratinocytes. (**a**) Principal component (PC) analysis of healthy donor, primary human KCs cultured for 24 hours with the indicated activated blood- or skin-derived T cell supernatants, compared with matched unstimulated controls. (**b**) PC analysis showing each blood (n=7) and skin (n=5) healthy donor-treated KC sample. (**c,d**) *Top*: Heat map showing z-score expression changes of KC genes (*rows*) occurring in response to culture with the indicated donor T cell population supernatant (*columns*). *Bottom*: Heat map showing cytokine production for each donor T cell population, based on quantification in Figure 1. All measures are scaled by quantile. IL-5, IL-17a, and IL-17f are truncated at the 95% quantile due to extreme outliers (c) The top 20 differentially expressed KC genes are shown in response to each indicated donor T cell population. (d) Expression of 18 well-characterized inflammatory and proliferative response elements in KC averaged across all donor samples for each T cell subset. (**e**) Dot plot showing functional enrichment analysis of differentially expressed, T cell-induced KC genes within the indicated modules (n=5-6 donors per population). The −log10 adjusted P value is plotted as a statistical measure for enrichment within each module.

To understand the biological significance of T cell-induced KC transcriptional states, we performed a functional enrichment analysis. Interferon response was largely T_h_1 cell-dependent while IL-10 immunoregulatory signaling and chemokine receptor pathways were highly enriched in response to T_h_17 cell supernatants (**Figure 2e, S2a-e**). Skin-derived T cell effects showed enrichment of keratinization and cell cycle pathway genes for CD103^neg^ and CD103^+^ fractions, respectively (**Figure 2d, 2e, S2f, S2g**). T cell activity broadly altered the baseline cytokine response potential of KCs, as evidenced by dramatic changes in their cytokine receptor expression (**Figure S3**). At rest, KCs constitutively expressed genes required for responsiveness to most T cell cytokines we measured, although CSF2RA (GM-CSF receptor) expression was near the lower limit of detection and neither IL2RA nor IL5R was detected. In response to stimulation, expression of IL10RB and IL22R1 were most highly induced in KCs by T_h_1 cells, corresponding to the level of IFNγ measured in stimulated cell supernatants. This is consistent with IFNγ induction of IL-10R as described in gut epithelial cells during barrier restoration ^22^. IL17RA and IL17RC expression by KCs, required for IL-17 signaling, were strongly induced by most CD4^+^CLA^+^ T cells tested. Expression of IL2RG, the common gamma chain, was strongly induced in response to stimulation, which potentiates the response to multiple other cytokines (IL-2, IL-4, IL-7, IL-9, IL-15). In these ways, CD4^+^CLA^+^ T cell subset activity can shape the outcome of cutaneous immune responses.

As observed in KCs, stimulation with blood and skin T cell supernatants had a strong effect on the transcriptional response in healthy donor dermal Fibs (**Figure 3a**). The largest effect size comes from T cell tissue of origin – blood versus skin – and while the impact of individual T_h_ populations was evident, this was less pronounced than that observed in KCs (**Figure 3a-c, S4**). The top DE genes in Fibs were separated into three main blocks: those associated with T_h_1 activity and high levels of the pro-inflammatory cytokine IFNγ; those associated with blood-derived T_h_ cells producing either type 2 or type 17-associated cytokines; and those associated with skin-derived T cells that produced proportionally more IL-22 and IL-9 than other subsets tested (**Figure 3c**). Each T cell population induced a different set of genes, though the main response in Fibs occurred largely downstream of either T_h_1/IFNγ or T_h_17/IL-17 (**Figure 3d, S4b, S4c**). Cell cycle and proliferation genes were elevated in response to skin-derived T cell supernatants, as observed in KCs, but also by T_h_17 and blood CD103^+^ T cell activity. A pathway analysis of the Fib response revealed broad engagement of cytokine signaling downstream of common CD4^+^ T cell cytokines including IFNγ, IL-4, IL-13, and IL-10 (**Figure 3e**). Skin CD103^neg^ T cells uniquely promoted metal ion and metallothionein signaling pathways, essential components of wound healing and the fibrotic response ^23^. As with KCs, treatment with T cell supernatants substantially altered cytokine receptor expression on Fibs (**Figure S5**).

**Figure 3.**
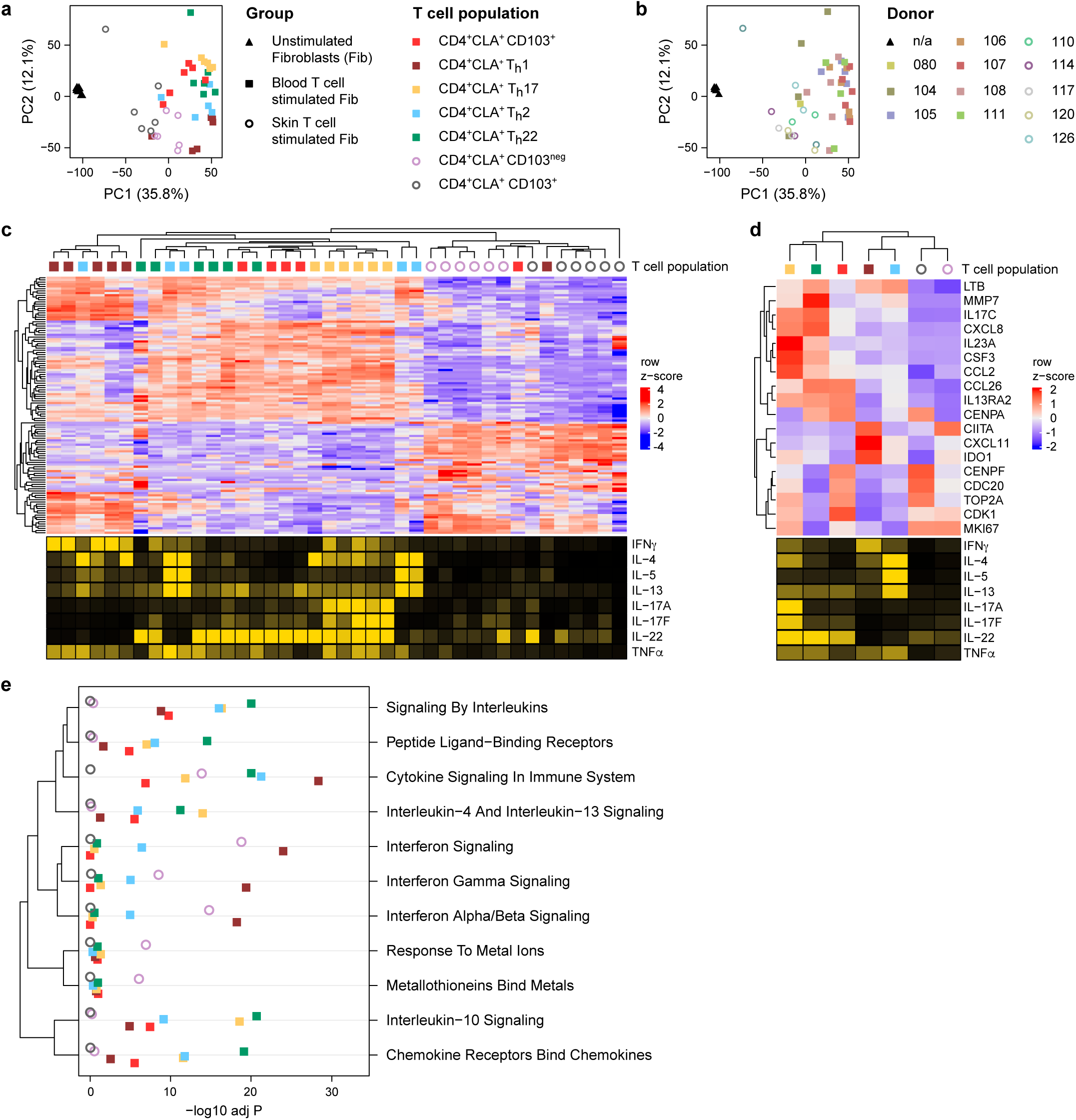
CD4^+^CLA^+^ T cells induce distinct transcriptional states in dermal fibroblasts. (**a**) Principal component (PC) analysis of healthy donor, primary human KCs cultured for 24 hours with the indicated activated blood- or skin-derived T cell supernatants, compared with matched unstimulated controls. (**b**) PC analysis showing each blood (n=7) and skin (n=5) healthy donor-treated Fib sample. (**c,d**) *Top*: Heat map showing z-score expression changes of Fib genes (*rows*) occurring in response to culture with the indicated donor T cell population supernatant (*columns*). *Bottom*: Heat map showing cytokine production for each donor T cell population, based on quantification in Figure 1. All measures are scaled by quantile. IL-5, IL-17a, and IL-17f are truncated at the 95% quantile due to extreme outliers. (c) The top 20 differentially expressed Fib genes are shown in response to each donor T cell population. (d) Expression of 18 well-characterized inflammatory and proliferative response elements in Fibs averaged across all donor samples for each T cell subset. (**e**) Dot plot showing functional enrichment analysis of differentially expressed, T cell-induced Fib genes within the indicated modules (n=5-6 donors per population). The −log10 adjusted P value is plotted as a statistical measure for enrichment within each module.

For example, induction of IFNGR1 and IL17RA in Fibs was induced by both blood- and skin-derived CD4^+^CLA^+^ T cells, consistent with our observation that a large part of the dermal Fib response to T cells is dependent on T_h_1/IFNγ and T_h_17/IL-17 signaling. In contrast, IL13RA2 expression is promoted by T_h_2, T_h_17, T_h_22 and blood CD103^+^ T cells, which each produce varying amounts of IL-13 and low levels of IFNγ. Once thought to be only a decoy receptor, IL13RA2 is an important mediator of IL-13-dependent fibrotic responses in barrier tissues ^24,25^. The baseline expression of IL31RA and OSMR were also elevated in stimulated Fibs, potentiating responsiveness to IL-31, a T cell cytokine involved in pruritis and atopic dermatitis ^26,27^. Finally, TGFBR1 and TGFBR2 expression were antagonized by blood- but not skin-derived CD4^+^CLA^+^ T cells. Collectively, these data show that CD4^+^CLA^+^ T cells can induce transcriptionally distinct states in epithelial KCs and dermal Fibs including those associated with inflammatory, regulatory, proliferative, and fibrotic gene programs of the skin.

### T cell-dependent gene signatures are enriched in the epidermis during inflammatory skin disease and normalized by anti-cytokine therapy

Phenotypically and functionally distinct T cell populations accumulate in lesional tissue during inflammatory and autoimmune skin diseases ^6,12^. Ps (∼3% prevalence in the US) and AD (∼7% prevalence in the US) are two common skin inflammatory diseases in which dysregulated T cell activity is central to the development or persistence of tissue pathology ^6,28,29^. Ps is primarily associated with elevated T_h_17 cell and IL-17 responses in the skin, whereas AD is canonically associated with T_h_2 cells and IL-13 activity, while IL-4 and IL-5 responses are mostly absent ^30,31^. The pathogenic role of T cell-derived cytokines in these diseases is highlighted by the success of biologic therapies: i.e. secukinumab and ixekizumab for Ps (anti-IL-17A); dupilumab and tralokinumab for AD (anti-IL4Rα, anti-IL13) ^32–35^. Using the structural cell gene signatures generated in Figures 2 and 3, we assessed the relationships between T cell activity, dysfunctional gene programs in skin structural cells during inflammatory disease, and patient responses to anti-cytokine therapy. As KCs are the majority population in the epidermis and alongside Fibs are the most abundant cells in human skin ^36^, we expected an analysis of full thickness skin biopsies would primarily reflect changes in KC and Fib gene programs.

We combined publicly available gene expression data of lesional, non-lesional, and healthy control donor skin from three clinical trials that demonstrated the efficacy of anti-IL-17A therapies in patients with Ps [GSE137218, GSE166388, GSE31652] ^37–40^, batch correcting to adjust for study-specific effects. The concatenated transcriptomic data sets clustered by disease state and treatment (**Figure 4a, S6a**). DE gene analysis revealed a pre-treatment lesional Ps skin gene expression profile that was altered over the course of anti-cytokine treatment, eventually resembling that of healthy control and non-lesional skin (**Figure 4a, 4b**), reflecting the known efficacy of these drugs.

**Figure 4.**
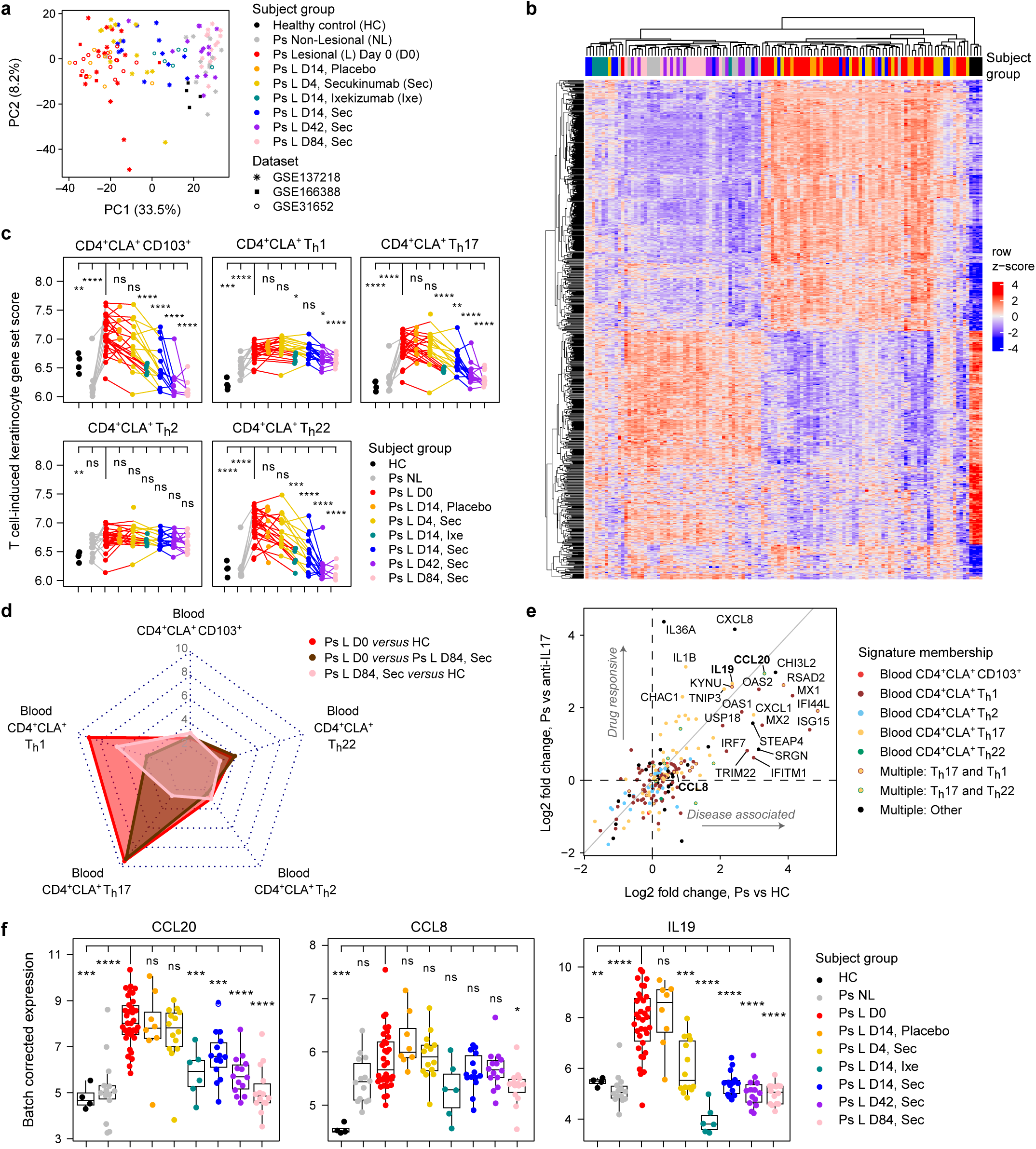
T cell-dependent gene signatures are enriched in the epidermis during psoriasis and normalized by anti-IL-17A. (**a**) PC analysis of gene expression for paired donor Ps lesional or non-lesional skin compared with healthy controls (HC) from the indicated public data sets. Subjects were treated with an anti-IL-17A therapy – secukinumab (Sec), ixekizumab (Ixe) – or placebo. Merged data sets were batch corrected to adjust for study-specific effects. (**b**) Heat map showing z-score expression changes of KC genes determined to be T cell-dependent (Figure 2) within public data, as shown in ‘a’ for the indicated groups. (**c**) Plots showing blood T cell-induced gene set scores, an averaged measure for the effect of each T cell subset on healthy donor KCs per subject group. (**d**) Radar plot showing T cell-induced KC signature gene enrichment values for the indicated pairwise comparisons from public data with −log10 p-values plotted along the radial axis. (**e**) Bivariate plot of the log2 fold change between subject groups for T cell-dependent KC genes (Ps vs HC, ‘Disease associated’; Ps vs anti-IL17A, ‘Drug responsive’). Individual genes are colored according to their T cell signature membership. (**f**) Plots showing batch corrected gene expression for each subject group for genes that have at least one expression-modulating Ps-associated SNP (i.e. eQTL). All statistical measures shown are compared to the Ps Lesional D0 group. Error bars indicate mean ± SD; ns = not significant, *p ≤ 0.05, **p ≤ 0.01, ***p ≤ 0.001, and ****p ≤ 0.0001 (Student’s *t*-test).

We next looked for T cell-dependent gene expression changes in Ps patient skin over the course of anti-IL-17A therapy. We found that gene signatures induced by T cell populations in KCs (**Figure 2**) were broadly increased as a part of the dysregulated transcriptional signature of disease (**Figure 4c**). The T_h_17-KC gene signature showed both the strongest association with the transcriptional dysregulation measured in Ps lesional skin and the subsequent response to therapy, compared to other T-dependent effects measured (**Figure 4d**). By day 84, both T_h_17-KC and T_h_22-KC signatures were significantly decreased in patient skin in response to therapy relative to day 0 lesional skin, consistent with the known role of IL-17 producing CD4^+^ T cells in Ps (**Figure 4c, 4d**). Of the 93 unique T_h_17-induced KC signature genes we identified (**Figure S2c**), 24 (26%) were DE between Ps and healthy skin samples. Of these 24 genes, 13 (54%) were significantly responsive to anti-IL-17A therapy while 11 (46%) were not (**Figure S6b**). We next assessed individual T cell-KC signature genes in lesional Ps skin compared either to healthy control or anti-IL-17A treated samples and found that many of the significant differences were T_h_17 cell-induced, disease-associated, and responsive to therapy (**Figure 4e**).

Further investigation revealed T_h_17-dependent KC genes that have Ps-associated GWAS SNPs near their transcriptional start site (TSS) (**Figure 4f**). The risk allele *rs7556897* is located 10.3kb from the CCL20 gene ^41^. It has a high variant-to-gene (V2G) score and impacts CCL20 as shown by expression and protein quantitative trait locus (eQTL, pQTL) mapping studies ^42–44^. Another, *rs9889296*, is located 75.9kb from CCL8 and is implicated by fine mapping as an eQTL ^41,45^. IL19 has 3 reported Ps GWAS hits: *rs3024493*, an eQTL for IL19 located 141bp from the TSS ^41^; *rs55705316*, located 10.6kb from the TSS with Hi-C data showing promoter proximity; ^46,47^ *rs12075255*, 17.5kb from TSS ^41^. CCL20 and CCL8 coordinate immune cell recruitment to the tissue ^48^ while IL19 is a part of the IL-17/IL-23 inflammatory axis of Ps and among the most DE cytokines measured between Ps ^49^. Our analysis demonstrated that CCL20, CCL8, and IL19 – genes elevated during the Ps epidermal inflammatory response and significantly impacted in response to anti-IL-17A therapy – are each uniquely induced by T_h_17-cells in KCs (**Figure 4f**). Published V2G mapping, eQTL and Hi-C studies complement our findings, reinforcing the functional relevance of these T cell-dependent target genes during Ps.

Not all the T-dependent KC genes we observed in lesional Ps skin were significantly impacted by anti-IL-17A therapy. Two examples are CXCL2, a neutrophil recruitment factor described to be part of the inflammatory axis of Ps ^50^, and NES, a gene encoding the intermediate filament nestin, which is expressed in skin epidermal stem cells and posited to be important during epithelial cell proliferation (**Figure S6b**) ^51^. CXCL2 and NES along with 9 other T_h_17-dependent genes are Ps-associated, but their expression is not significantly altered by anti-IL-17A blockade. Thus, we were able to discern a mixed effect of cytokine blockade on T cell-dependent gene expression in the skin, though it remains unclear which of these genes are central indicators of patient outcome to therapy.

We took a similar approach to analyze publicly available skin RNA-seq data from an AD clinical trial that tested the efficacy of IL-4R blockade by dupilumab [GSE157194] ^52^ and from matched AD patients and controls [GSE121212] ^53^. Most of the variance in the combined data set could be explained by disease state, while a smaller effect was evident in response to treatment (**Figure 5a, S7a**). We focused our analysis on DE genes that we identified as T cell-dependent in KCs (**Figure 2**). Using this strategy, the data separated largely by tissue type; lesional, non-lesional, and healthy skin. Samples in dupilumab treated groups were dispersed throughout hierarchical clusters, indicating a mild treatment effect on total T cell-dependent KC genes (**Figure 5b**). When comparing subset-specific T cell-KC gene expression across all subject groups we found a significant effect of dupilumab on the expression of CD4^+^CLA^+^ T_h_2, T_h_22, and CD103^+^ T cell-dependent genes within lesional skin (**Figure 5c**). All T cell-KC gene sets were significantly upregulated within lesional skin samples compared with those from healthy skin, an effect that was diminished in response to therapy (**Figure 5c, 5d**). Dupilumab targets IL-4 and IL-13 activity, two cytokines made co-produced in abundance by CD4^+^CLA^+^ T_h_2 cells (**Figure 1d**). We identified a set of seven uniquely T_h_2-induced KC genes that were significantly elevated in lesional AD skin compared to controls (**Figure 5e, S7b**). Of these, CA2, SLC5A5, and SLC26A9 were significantly reduced in lesional skin following dupilumab treatment (**Figure S7c**). CA2 encodes a carbonic anhydrase important for cellular pH and ion homeostasis, previously shown to be elevated in AD patient skin and differentiated KCs ^54,55^. SLC5A5 and SLC26A9 encode iodine and chloride channel proteins, respectively, and neither have a known role in promoting pathology during AD.

**Figure 5.**
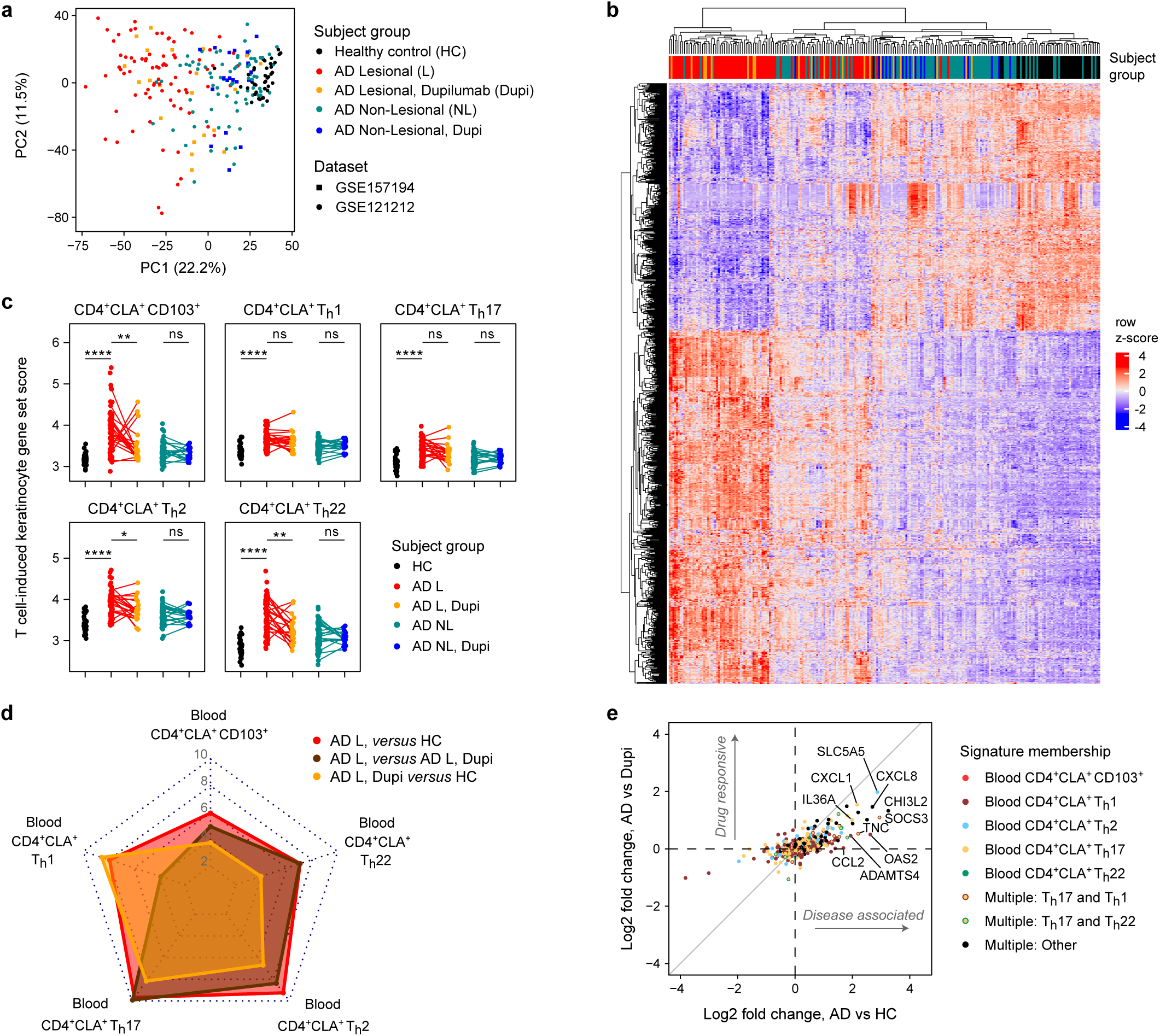
T cell-dependent gene signatures are enriched in the epidermis during atopic dermatitis and normalized by anti-IL-4Rα therapy. (**a**) PC analysis of gene expression for AD paired donor lesional and non-lesional skin compared with healthy controls (HC) from the indicated publicly available data sets. AD patients were treated with dupilumab (Dupi), an anti-IL-4Rα drug. Merged data sets were batch corrected to adjust for study-specific effects. (**b**) Heat map showing z-score expression changes of KC genes determined to be T cell-dependent (Figure 2) within public data, as shown in ‘a’ for the indicated groups. (**c**) Plots showing blood T cell-induced gene set scores, an averaged measure for the effect of each T cell subset on healthy donor KCs per subject group. (**d**) Radar plot showing T cell-induced KC signature gene enrichment values across the indicated pairwise comparisons from public clinical trial data (AD and HC) with −log10 p-values plotted along the radial axis. (**e**) Bivariate plot of the log2 fold change for T cell-dependent KC genes within public clinical trial data from AD or HC subjects (AD vs HC, ‘Disease associated’; AD vs anti-IL17A, ‘Drug responsive’). Individual genes are colored according to their T cell signature membership. Error bars indicate mean ± SD; ns = not significant, *p ≤ 0.05, **p ≤ 0.01, ***p ≤ 0.001, and ****p ≤ 0.0001 (Student’s *t*-test).

Our findings demonstrate that T cell-dependent KC gene networks experimentally derived *in vitro*: (1) are enriched *in vivo* within human skin; (2) are associated with cutaneous inflammatory disease; and (3) some but not all are normalized in response to anti-cytokine biologic therapy. Further disease relevance is highlighted by fine-mapping studies that link GWAS-identified SNPs to the expression of T-dependent genes we describe. Thus, we validate a method of identifying candidate genes to explain immune-dependent effects during inflammatory disease and to deconvolute the patient response to therapy.

### Skin CLA^+^ T cells and the gene signatures they induce in dermal fibroblasts are present in healthy individuals and reduced during inflammatory skin disease

T cell infiltration and aberrant cytokine production cause fibrosis and vascular abnormalities in people with inflammatory disease ^56^. SSc is a rare inflammatory condition (∼0.1% prevalence in the US) with the highest mortality of any rheumatic illness. In the skin, pathology is associated with elevated T_h_2 cell infiltration, myofibroblast and alternatively activated macrophage differentiation ^57,58^. Effective therapies for SSc are severely lacking and ongoing clinical trials are targeting T cell activation and cytokine signaling networks (i.e. IL-4, IL-13, TGFβ, IL-6, JAK/STAT) ^59^. Dermal tissue pathology and fibroblast dysfunction has also been described during Ps and is connected to T cell cytokine activity (i.e. IL-17, IL-23, TNFα, IL-36) ^60,61^. Despite evidence implicating T cell activity in fibroblast dysregulation, the critical T cell-Fib interactions of healthy and diseased skin have not been established.

To determine whether and to what extent T cell-dependent gene signatures can be found in dermal Fib populations and whether this changes during disease, we analyzed publicly available scRNA-seq data from SSc patients and healthy controls [GSE195452] ^3^. In this study, the authors identified 10 transcriptionally distinct Fib subsets including Fib-LGR5, a population that is selectively enriched in healthy skin and significantly diminished with the progression of SSc. The unique functions of these Fib populations and their contribution to disease remain undefined. Our analysis of cells annotated ‘fibroblasts’ from GSE195452 confirms that the Fib-LGR5 population is preferentially found in healthy skin compared with SSc samples (**Figure 6a, 6b**) and the fraction of Fib-LGR5 is diminished progressively in patients with limited and diffuse disease (**Figure S8a, S8b**). The Fib-MYOC2 and Fib-POSTN subsets were reciprocally elevated in the skin of SSc patients, which correlated with higher autoantibody levels and an increase in skin disease score (**Figure S8b, S8c**). T cell-dependent gene signatures (**Figure 3**) were detected throughout newly described Fib subsets in pooled data from healthy control and SSc subjects. The most pronounced effect observed was the enrichment of skin CD4^+^CLA^+^ T cell-induced genes in the Fib-LGR5 population (**Figure 6c**). The skin CD4^+^CLA^+^ CD103^+^ T cell-dependent Fib gene signature is specifically elevated in the fraction of Fib-LGR5 cells of healthy donors and reduced in diffuse SSc subject skin while the opposite pattern was seen for the Fib-MYOC2 subset (**Figure 6d**).

**Figure 6.**
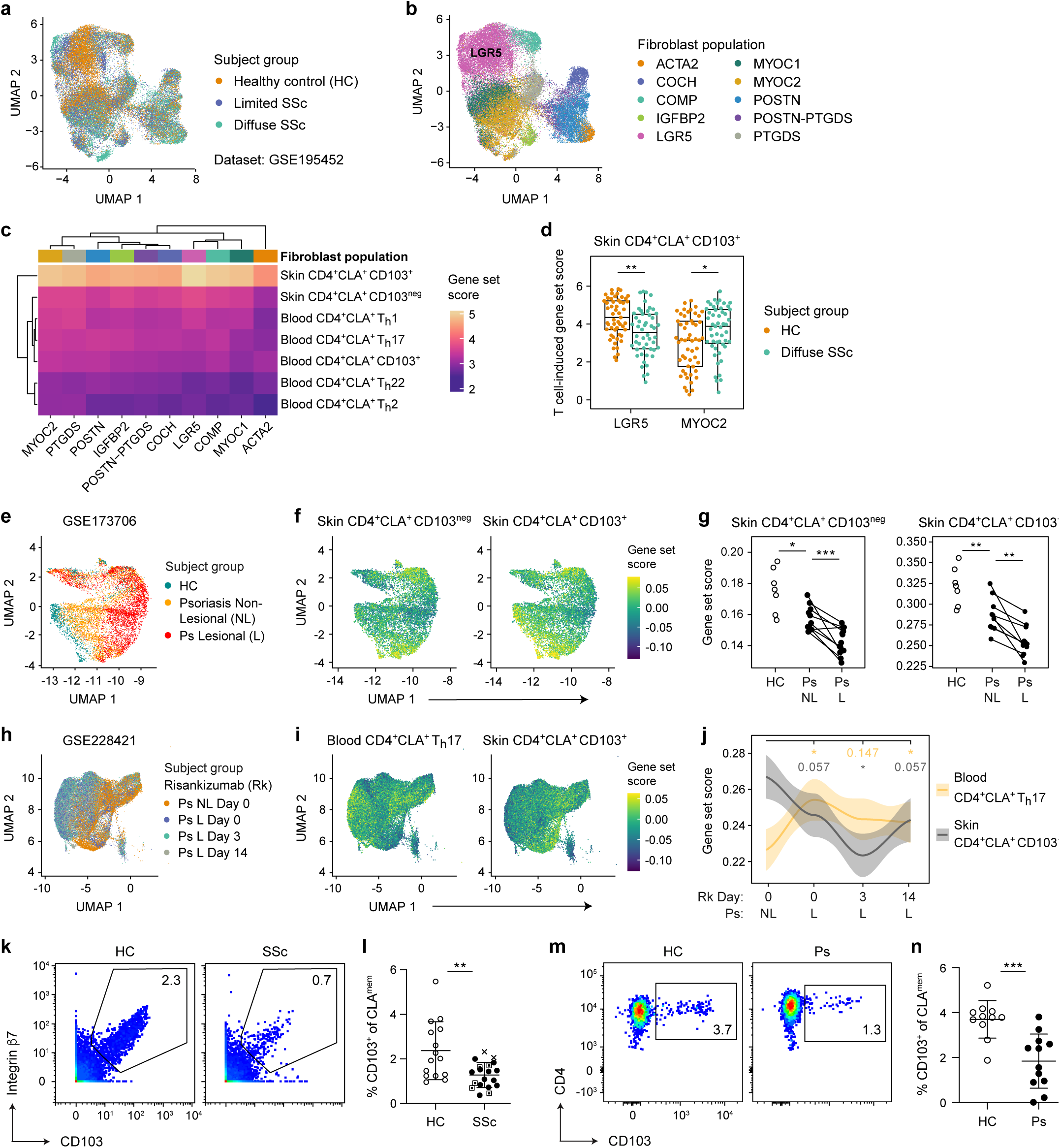
Skin T cell-induced gene signatures and CD4^+^CLA^+^ CD103^+^ T cells are elevated in healthy subjects and reduced during inflammatory skin disease. (**a-j**) T cell-induced Fib gene sets (Figure 3) were used to analyze (a-d) public scleroderma and (e-j) psoriasis skin scRNA-seq data, filtered on cells annotated as ‘fibroblasts’. Ps lesional and non-lesional skin is from paired donors. (**a-d**) Uniform Manifold Approximation and Projection (UMAP) dimensionality reduction plots showing assignment by (a) subject group and (b) defined Fib subset [GSE195452]. (c) Heat map showing the enrichment of T cell-dependent gene signatures for each Fib population pooled across all subject groups. (d) Plot showing T cell-dependent gene signature enrichment in Fib subsets, pseudobulked at the subject level. (**e-g**) UMAP plots showing assignment (e) by subject group and (f) skin T cell-dependent gene set enrichment, (g) quantified per group [GSE173706]. (h-j) UMAP plots of patient data [GSE228421] before (Day 0) and after (Days 3, 14) IL-23 blockade (Risankizumab, Rk) showing assignment (h) by subject group and (i) T cell-dependent gene set enrichment, (j) quantified per group. (**k-m**) Cytometric analysis of blood CD4^+^CLA^+^ CD103^+^ T cells. (k) Representative fluorescence cytometry plots and (l) quantification of T cells in HC (n=15) or SSc (n=19) patient blood, gated on CD103, integrin β7, and pre-gated on CLA^mem^, as in Figure 1. SSc samples: circle = limited disease, square = diffuse, X = unclassified. (m) Representative mass cytometry plots and (n) quantification of T cells in HC (n=11) or Ps (n=12) patient blood, gated on CLA^mem^, as in Figure S8. Error bars indicate mean ± SD; ns = not significant, *p ≤ 0.05, **p ≤ 0.01, ***p ≤ 0.001, and ****p ≤ 0.0001 (d, Student’s *t*-test; l,n, Mann-Whitney U).

We next sought to determine whether transcriptional effects induced by CD4^+^CLA^+^ T cell subsets might contribute to the Ps inflammatory fibroblast axis. Gene expression analysis of skin fibroblasts [GSE173706] ^61^ showed subject-level distinction between healthy control, non-lesional, and lesional groups (**Figure 6e, S8a**). We observed that both skin CD4^+^CLA^+^ CD103^neg^ and CD103^+^-induced gene sets (**Figure 3**) are preferentially found in healthy control skin Fibs compared to non-lesional and lesional Ps (**Figure 6f, 6g**). We saw no significant effects when measuring blood CD4^+^CLA^+^-induced Fib genes (**Figure S8d**). We similarly found skin CD4^+^CLA^+^ CD103^+^ T cell-induced gene programs are reduced in lesional Ps skin compared with paired donor non-lesional biopsies (**Figure 6h-6j, S8a**) in a second Ps clinical cohort [GSE228421] ^60^. This study assessed the IL-23 biologic Risankizumab in five Ps subjects, describing dermal fibroblasts gene expression changes occurring over a brief 2-week period after therapy. Consistent with the known role of IL-23 as an activator of the IL-17 signaling cascade, we observed an increase in blood CD4^+^CLA^+^ T_h_17-induced signature in fibroblasts from lesional tissue. IL-23 blockade promoted a reversal of the Th17-KC signature back to baseline, but the effect was mixed at the two early timepoints post treatment (**Figure 6j**). We also detected a modest enrichment of blood CD4^+^CLA^+^ CD103^+^ T cell-induced genes in the lesional Ps group (**Figure S8e**).

CD4^+^CLA^+^ CD103^+^ T cells in the blood represent a migratory fraction of T_RM_ that express a transcriptional and cytokine profile indicative of a role in wound healing and tissue repair ^11^. As CD4^+^CLA^+^ CD103^+^ T cell-induced gene signatures are altered between healthy control and diseased subject skin, we sought to determine if T cell abundance was also impacted. We measured the frequency of CD4^+^CLA^+^ CD103^+^ T cells in the blood of SSc and Ps patients and found a reduction in both disease groups with respect to matched healthy controls (**Figure 6k-6n, S8f**), suggesting that the regulation of the CD103^+^ T cell population is important during inflammatory skin disease. Thus, we show that skin CD4^+^CLA^+^ T cell activity supports the transcriptional programs of healthy skin-associated Fibs and that CD103^+^ T cells in circulation are diminished during inflammatory skin disease.

## DISCUSSION

In this study, we constructed an atlas of T cell-induced gene states in human skin – a catalog of epidermal keratinocyte and dermal fibroblast transcriptional reprogramming by different CD4^+^CLA^+^ T_h_ cell populations – then applied our findings towards a deeper understanding of clinical data sets. Although the use of CLA to track cutaneous T cells goes back nearly three decades, ^8,9^ our approach uniquely combines CLA identification of both blood- and skin-derived T_h_ cells together with chemokine receptor gating strategies and CD103 designation of migratory T_RM_ ^10,11^ to isolate broad and functionally diverse CD4^+^CLA^+^ T_h_ cells, which we leverage to study the changing transcriptional landscape of healthy and inflamed human skin.

T cells elicited significant effects on both primary KCs and Fibs, substantially altering the transcriptional landscape of each cell type. Cytokine-dependent gene modules were broadly enriched, especially through the activity of blood-derived CLA^+^ T cells. This included expected gene set enrichment such as CLA^+^ T_h_1-mediated IFN-associated gene expression, and CLA^+^ T_h_2-mediated IL-4 and IL-13 responses in both KCs and Fibs. Skin-derived T cells were instead more likely to promote gene networks required to execute critical homeostatic functions of skin. Keratinization, cornification, and proliferation processes – essential for maintenance and renewal of a healthy epithelium ^62^ – were induced specifically in KCs by CD103^neg^ or CD103^+^ skin T cells, while metal ion response pathways involved in wound healing ^23^ were induced by CD103^neg^ T cells, which was only observed in Fibs. Our findings demonstrate the broad functional capacity of each these T cell populations and suggest how changes in the CD4^+^CLA^+^ pool would impact both the inflammatory response against harmful pathogens and the upkeep of homeostatic tissue functions such as barrier maintenance, recovery from injury, and regulated self-renewal.

Blocking T cell cytokine responses during Ps (i.e. IL-17) and AD (i.e. IL-4/IL-13) reduces leukocyte accumulation in the tissue and reverses skin pathology. While many studies have shown the efficacy of these drugs, not all patients meet the desired clinical endpoints and severe adverse events occur in a small but significant proportion of individuals, warranting further investigation to define the mechanisms of drug action and to identify new candidate drugs as alternative therapies ^63–68^. TNFα or IL-23 blocking therapies, for example, offer substitutes for Ps patients failing anti-IL-17A therapy, and the IL-13 specific agonist tralokinumab is an alternative to dupilumab in AD. Establishing biomarkers of disease and the response to therapy will help guide the most effective course of treatment on an individual patient level. Our study adds important context that can help interpret clinical outcomes following anti-cytokine therapy. We describe the T cell-dependent gene networks in KCs that are enriched in lesional skin of Ps and AD patients compared with non-lesional and healthy controls, and we demonstrate how the expression of specific T-dependent genes are impacted during anti-cytokine therapy in patient skin.

By combining this approach with public immunogenetics data from Ps patients, we identified CCL20, CCL8, and IL19 as T_h_17 cell-induced targets of anti-IL-17A therapy. Expression of each of these genes is elevated in lesional skin compared with controls, returned to baseline following therapy, and each contains GWAS risk alleles for Ps that are eQTL. Taken together with published studies, our findings suggest these genes are part of a cooperative functional network that promotes Ps: CCL20 recruits IL-17 producing disease-associated CCR6^+^ T_h_17 cells to the skin; ^48,69^ CCL8 recruits CCR5^+^ CD4^+^ T cells that are involved in barrier integrity maintenance and implicated in Ps pathogenesis; ^70,71^ IL-19 expression is positively associated with Psoriasis Area and Severity Index – a measure of skin pathology. Expression of IL-19 is both IL-17-dependent and further enhances the effects of IL-17A, increasing IL-23p19 expression and production of CCL20 ^49^. IL-19 is part of the IL-23/IL-17 inflammatory signaling network of Ps and is a proposed biomarker of disease. Guselkumab (anti-IL-23) is an approved drug for Ps that showed superior long-term efficacy to secukinumab, ^72^ but no current therapy targets other components of the T cell/KC/Ps axis we describe such as IL-19, CCL8, or skin-tropic CCR5^+^ T cells.

We also describe genes that are uniquely induced by CLA^+^ T_h_2 cells in KCs, which are significantly altered in lesional AD patient skin and responsive to dupilumab. For example, CA2 was previously shown to be induced in KCs and engineered skin equivalents by IL-4 and IL-13 ^55,73^. In comparison NTRK1, an early IL-13 target involved in allergic responses, is not clearly connected to AD ^74^, and our findings suggest a stronger link. Other genes we identify, such as TNC, CHI3L2 and SLC5A5, were shown to be upregulated in AD patient lesional skin, ^75–77^ but further research is required to determine the relevance of these factors in disease and the response to treatment. Importantly, some *but not all* T-dependent KC genes enriched in lesional patient skin are responsive to biologic therapy, which could indicate the presence of stable epigenetic changes imprinted during disease. Our study establishes a strategy to assess the transcriptional response to biologics, which could help identify novel cellular and genetic targets for future therapeutic intervention.

Our analyses of skin Fib responses suggests that a balance of functionally distinct CLA^+^ T cell populations is critical to Ps and SSc disease progression. We show across multiple patient cohorts and diseases that skin CD4^+^CLA^+^ CD103^+^ T cells support the gene programs of healthy dermal populations such as Fib-LGR5 that are significantly reduced during SSc. This raises the possibility that CD4^+^CLA^+^ CD103^+^ T cell activity is directly required to maintain Fib-LGR5 and that loss or dysregulation of these T cells contributes to the development of myofibroblasts known to drive fibrosis such as Fib-MYOC2, -SFRP2, or -PRSS23 ^5,78^. The LGR5 expressing population is transcriptionally similar to Fib-PI16 described previously, ^2^ with both Fib-LGR5 and -PI16 independently shown to be reduced in SSc patient skin ^3,79^. Individuals with severe SSc and reduced Fib-LGR5 are the most likely to have anti-topoisomerase I (scl-70) autoantibodies, which puts them at the highest risk for severe pulmonary fibrosis and end-organ disease ^80^, reinforcing the importance of understanding immune-based regulation of healthy skin-associated Fibs.

Previously, we published that skin CD4^+^CLA^+^ CD103^+^ T_RM_ cells uniquely co-produce IL-13 and IL-22 and have a TGFβ gene signature, implicating them in tissue repair, barrier maintenance, and homeostatic function ^11^. We now report that the migratory fraction of CD4^+^CLA^+^ CD103^+^ T cells in the blood is reduced in both SSc and Ps patients, supporting the notion that these cells are important to disease development. Whether these T cells are also altered in the skin, if they become dysfunctional during disease, and if they directly impact the activity or survival of disease-relevant populations in the skin such as Fib-LGR5 remain open, yet exciting questions. A therapeutic strategy focused on reinforcing the function or number or healthy skin-associated T cell and Fib populations could be a useful adjunct to anti-cytokine therapies that are aimed at minimizing the impact of deleterious immune cells.

Our study design and subsequent analysis contain some caveats and limitations. First, we measure 13 analytes produced by activated T cells, which accounts for only a fraction of the cytokines, metabolites and other soluble factors that are part of the T cell-dependent regulation of skin structural cell gene programs. Second, our study uses KCs and Fibs from a single anatomical skin site and donor. Thus, we are unable to comment on site-specific differences, for example owing to changes in moisture content, microbiome, or hair follicle density across this large barrier tissue, or population level variability in T cell-induced responses. Finally, the data from several publicly available interventional clinical trials that we assessed in this study were generated using either bulk sequencing or microarray methods, and some were smaller studies with limited power. Additionally, not all data sets we assessed showed a significant enrichment of T cell-dependent gene programs (*data not shown*) ^2,5,79,81^. Future analyses are required to validate the essential T cell- and cytokine-directed gene networks in KC or Fib subsets during health and disease across this large barrier tissue.

The atlas of T cell-dependent gene expression responses that we introduce in this study represents a new tool to facilitate a deeper understanding of the complex biology and transcriptional networks of skin. Using our data to deconvolute published clinical studies has yielded new insights into immune-directed pathogenesis of Ps, AD, and SSc, and demonstrates the importance of T cells in the upkeep of structural cell subsets in healthy skin. Thus, our study presents and validates a broadly applicable, systems-based approach to dissecting the immune-dependent contributions to tissue homeostasis, inflammation, and fibrosis.

## MATERIALS AND METHODS

### Human subject enrollment and study design

The objective of this research was to characterize the effect of human T cells on the gene expression profile of epithelial KCs and dermal Fibs. Immune cell isolation was performed using healthy blood and skin tissue – donor range 32 to 71 years of age. Human skin was obtained from patients undergoing elective surgery – panniculectomy or abdominoplasty. The cohort size was selected to ensure a greater than 80% probability of identifying an effect of >20% in measured variables. All samples (**Table S5**) were obtained upon written informed consent at Virginia Mason Franciscan Health (Seattle, WA, USA) and the Dr. D. Y. Patil Medical College, Hospital, and Research Centre (Pune, India). All study protocols were conducted according to Declaration of Helsinki principles and approved by the Institutional Review Board of Benaroya Research Institute (Seattle, WA, USA), the Institutional Human Ethics Committees of Dr. D. Y. Patil Medical College, Hospital, and Research Centre (Pune, India) and CSIR – Institute of Genomics and Integrative Biology (New Delhi, India).

### Sex as a biological variable

Our study examined both male and female tissue with similar findings reported for both sexes. The study was not powered to detect sex-based differences.

### Isolation of T cells from blood

PBMC were isolated using Ficoll-Hypaque (GE-Healthcare; GE17-1440-02) gradient separation. T cells were enriched using CD4 microbeads (Miltenyi; 130-045-101) then rested overnight at a concentration of 2 x 10^6^ cells/mL in ImmunoCult™-XF T Cell Expansion Medium (StemCell; 10981) with 1% penicillin/streptomycin (Sigma-Aldrich; P0781) in a 15 cm^2^ dish. The next morning cells were harvested and prepared for cell sorting, as indicated.

### Isolation of T cells from skin

Fresh surgical discards of abdominal skin were treated with PBS + 0.1% Primocin (Invitrogen; ant-pm-1) for 5 minutes to eliminate contaminating microorganisms. Sterile tissue processing was performed on ice and skin was periodically sprayed with 1x PBS to keep samples from drying out. A biopsy tool (Integra™ Miltex™) was used to excise 4mm tissue biopsies from each sample. The subcutaneous fat was removed from biopsies before an overnight digestion of approximately 16 hours at 37°C, 5%CO_2_. The digestion mix contained 0.8 mg/mL Collagenase Type 4 (Worthington; LS004186) and 0.04 mg/mL DNase (Sigma-Aldrich; DN25) in RPMI supplemented with 10% pooled male human serum (Sigma-Aldrich; H4522, lot SLC0690), 2 mM L-glutamine, 100 U/mL penicillin, and 100 mg/mL streptomycin. We used 1 mL of RPMI digestion media for each 40-50mg tissue in a 6-well non-tissue culture treated dish. The next morning tissue samples were washed excess RPMI + 3% FBS and combined into a single tube per donor. Filtration on a 100μM membrane was performed at each wash step, five total on average, to generate a single-cell suspension of skin mononuclear cells free of contaminating fat and debris. Skin mononuclear cell suspensions were then prepared for cell sorting, as indicated.

### Cytometry and cell sorting

*Fluorescence cytometry* – Cell labelling of surface antigens on blood- and skin-derived T cells was performed using fluorescently tagged antibodies diluted in cell staining buffer containing HBSS and 0.3% BSA (**Table S6**). Viability dye staining using Fixable LiveDead was performed first in HBSS containing no additional protein, as recommended by the manufacturer. Surface staining was performed at 5% CO_2_, 37°C for 20 minutes. Labeled cells were then either acquired on a BD LSR™ Fortessa cell analyzer or prepared for cell sorting.

#### Cell sorting

Fluorescently labelled samples were placed in cell sorting buffer – for yield sorts, HBSS + 5.0% BSA, for purity sorts HBSS + 1.0% BSA – using a BD FACSAria™ Fusion (85μM nozzle, 45 p.s.i.). The indicated conventional T_h_ cell populations (T_Conv_) were sorted from blood and T_Reg_ were excluded by purity sort following CD4^+^ T cell microbead enrichment. Skin mononuclear preparations were first enriched by a CD3 yield sort followed by purity sort of CD4^+^ T_Conv_ and exclusion of T_Reg_. Additional sample collection was performed using the BD LSR™ Fortessa cell analyzer.

#### Mass cytometry

Frozen PBMC samples were prepared as described, then rested overnight in ImmunoCult™-XF T Cell Expansion Medium supplemented with 1% penicillin/streptomycin and 5% human serum. Human pan T cell isolation was performed (Miltenyi; 130-096-535) and 0.5 x 10^6^ to 2 x 10^6^ purified T cells were stained for analysis, as follows. Cell viability staining using 5 µM Cisplatin (Enzo Life Sciences; ALX-400-040) in PBS. After 1 minute at room temperature the reaction was quenched. Metal-tagged antigen labeling using antibody cocktail (**Table S6**) diluted in Maxpar® Cell Staining Buffer (Standard BioTools; 201068) for 30 minutes at room temperature. Labeled cells were resuspended with Maxpar Intercalator-Ir intact stain in Fix and Perm Buffer at 4°C, overnight (Fluidigm; 201192A and 201067). Cells were resuspended in ultrapure water containing 1/5^th^ volume EQ Four Element Calibration Beads (Standard BioTools; 201078) and acquired on a Standard BioTools CyTOF Helios at an event rate of 300-400 cells per second. Samples were processed in four subsequent batches over the course of one week. All fluorescence and mass cytometry data (.FCS 3.0 or .FCS 3.1) were analyzed using FlowJo software (BD, v.10.9).

### T cell stimulation and cytometric quantification of cytokines

Sorted T cell populations were pelleted by centrifugation at 300 x g for 5 minutes and resuspend in ImmunoCult™-XF T Cell Expansion Medium with 1% penicillin/streptomycin. Cultured T cells were stimulated for 48 hours at 1 x 10^6^ cells/mL with ImmunoCult™ Human αCD3/αCD28/αCD2 T Cell Activator reagent (StemCell; 10970) used at the manufacturer recommended concentration. Cytokine containing supernatants from each unique donor were harvested and stored at −20C°. Frozen blood and skin activated T cell supernatants from all donors were thawed as a single batch to assess cytokine concentrations by cytometric bead array (BioLegend; LEGENDplex™ reagent) using a custom analyte panel: GM-CSF, IFNγ, IL-2, IL-4, IL-5, IL-6, IL-9, IL-10, IL-13, IL-17A, IL-17F, IL-22, TNFα. Data were analyzed using the manufacturer provided LEGENDplex software. The remaining thawed T cell supernatants were used immediately, without additional freeze-thaws, in culture of KCs or Fibs, as described. Unstimulated controls were generated using a single freeze-thawed aliquot of cell-free ImmunoCult™-XF T Cell Expansion Medium.

### Keratinocyte and fibroblast cell culture and activation

Paired primary human epithelial KCs and dermal Fibs from the eyelid of a single healthy donor were purchased from a commercial vendor (ZenBio; KR-F or DF-F) and cultured at low passages (p = 2-7) using either the KGM™ Keratinocyte Growth Medium BulletKit™ (Lonza; CC-3111) for KCs or DMEM supplemented with 10% FBS, 2 mM L-glutamine, 100 U/ml penicillin, and 100 mg/ml streptomycin (Gibco™; 10938025) for Fibs. Cytokine containing T cell supernatants were diluted using a 1:1 ratio of appropriate cell culture media and added to KCs or Fibs growing at 50% confluence – approximately 1-2 x 10^4^ – in a 96-well flat bottom plate. Structural cells were cultured with activated T cell supernatants for 24 hours prior to isolation for RNA-sequencing (RNA-seq). Unstimulated controls received a 1:1 ratio of fresh T cell media combined with either KC or Fib culture media.

### Bulk RNA-sequencing (RNA-seq)

High-quality total RNA was isolated from approximately 2 x 10^4^ T cell supernatant-treated KCs or Fibs using TRIzol^TM^ Reagent (Invitrogen^TM^; 15596026). Next, cDNA was prepared using the SMART-Seq v4 Ultra Low Input RNA Kit for Sequencing (Takara). Library construction was performed using the NexteraXT DNA sample preparation kit (Illumina) using half the recommended volumes and reagents. Dual-index, single-read sequencing of pooled libraries was run on a HiSeq2500 sequencer (Illumina) with 58-base reads and a target depth of 5 million reads per sample. Base-calling and demultiplexing were performed automatically on BaseSpace (Illumina) to generate FASTQ files.

### Analysis of bulk RNA-seq data

The FASTQ files were processed to remove reads of zero length (fastq_trimmer v.1.0.0), remove adapter sequences (fastqmcf tool v.1.1.2), and perform quality trimming from both ends until a minimum base quality ≥ 30 (FASTQ quality trimmer tool v.1.0.0). Reads were aligned to the human reference genome (build hg38) with TopHat (v.1.4.0) and read counts per Ensembl gene ID were quantified with htseq-count (v.0.4.1). Quality metrics for the FASTQ and BAM/SAM files were generated with FastQC (v.0.11.3) and Picard (v.1.128). Processing of FASTQ and BAM/SAM files was executed on the Galaxy workflow platform of Globus genomics. Statistical analysis of gene expression was assessed in the R environment (v.4.3.2). Sample inclusion for final analysis was based on a set of pre-established quality control criteria: total number of fastq reads > 1 x 10^6^; mapped reads > 70%; median CV coverage < 0.85. Randomization was not applicable as no treatment or intervention groups were included in the study. Blinding was not applicable as no treatment groups were compared.

### Additional bioinformatic analyses

All differential expression and pathway enrichment analyses were performed in R. To assess differential expression, the limma package (v.3.58.1) was used ^82^. A log2 expression fold change of at least 1 in magnitude and an FDR of less than 0.05 were used as cutoffs to define differentially expressed genes. Individual blood- and skin-T cell subset induced gene sets were formed by first filtering out any differentially expressed genes shared across all treatment groups – broadly excluding non-specific T cell effects on keratinocytes or fibroblasts – and then selecting up to the top 200 differentially upregulated genes ranked by FDR and associated with a T cell subset stimulation condition.

In analysis of public bulk microarray and RNA-seq GEO datasets, the ComBat function from the sva package (v.3.50.0) was used to adjust for dataset-specific batch effects ^83^. Pathway enrichment analysis was performed using the enrichR package (v.3.2) ^84^ with the Reactome 2022 database ^85^ to query pathways associated with T cell-induced genes. T cell signatures were treated as a custom database and expression of these signatures were examined as lists of differentially expressed genes associated with Ps or AD.

GWAS hits associated with the trait “psoriasis” were downloaded from the NHGRI-EBI GWAS Catalog on February 7^th^, 2024. Within these results, GWAS SNPs with a “DISEASE/TRAIT” value of “Generalized pustular psoriasis”, “COVID-19 or psoriasis (trans-disease meta-analysis)”, or “Paradoxical eczema in biologic-treated plaque psoriasis” were excluded to ensure Ps-specific SNPs were analyzed ^86^.

Single cell gene expression data from SSc and Ps cohort studies was re-analyzed using the Seurat R package (v.4) ^87^. CellTypist was used to identify fibroblasts using an adult human skin reference model ^88–90^. For SSc public data, pseudobulking was applied to defined SSc Fib clusters [GSE195452] ^3^. The average expression of T cell-induced genes was then computed for each pseudobulk profile as a gene set score. For Ps public data, pseudobulking was applied to tissue type or subject/visit group as indicated [GSE173706] ^61^ and [GSE228421] ^60^. The average expression of T cell-induced genes was then computed for each pseudobulk profile as a gene set score. Gene set scores for individual cells were computed using the AddModuleScores() method within Seurat ^91^.

## Supporting information

Supporting Data Values

## DATA AVAILABILITY STATEMENT

Values for all data points presented in figure graphs are reported in the **Supporting Data Values** file. All gene expression data sets generated in this study are available to the public in the Gene Expression Omnibus, **GSE272623**. Computer code generated and used for analysis in this study is available in Github: Commit URL, https://github.com/BenaroyaResearch/Morawski-T-cell-transcriptional-programs-in-skin; Commit ID, ab964c4.

## CONFLICT OF INTEREST STATEMENT

The authors declare no competing interests.

## ACKNOWLEDGEMENTS

We are grateful to: Kassidy Benoscek-Narag, Sylvia Posso, Thien-Son Nguyen, and the Clinical Research Center coordinators in the Benaroya Research Institute (BRI) Center for Interventional Immunology for human subject recruitment, sample collection and screening; the BRI Genomics, Flow Cytometry, and Human Immunophenotyping facilities for technical assistance and expertise; the BRI Scientific Writing Group for critical reading of the text. BRI cytometry and genomics equipment used in this study were generously supported by the M.J. Murdock Charitable Trust. This work was supported by a New Investigator research award (National Scleroderma Foundation) and R21AI185642 (NIAID, NIH) to PAM, R01AI127726 to IKG and DJC (NIAID, NIH), and R01AI169893 to HAD, IKG and DJC (NIAID, NIH).

## AUTHOR CONTRIBUTIONS (CrediT STATEMENT)

Conceptualization: HAD, IKG, JSC, DJC, PAM; Data Curation: HAD; Formal Analysis: HAD, DJC, PAM; Funding Acquisition: HAD, IKG, DJC, PAM; Investigation: HAD, MLF, SRV, AS, PAM; Methodology: HAD, DJC, PAM; Project administration: PAM; Software: HAD; Resources: JDS, WPS, AG, AS, JSC; Supervision: IKG, DJC, PAM; Validation: HAD, PAM; Visualization: HAD, DJC, PAM; Writing – original draft: HAD, PAM; Writing – review and editing: HAD, IKG, DJC, PAM.

## SUPPLEMENTARY MATERIAL

**Document S1.** Supporting data values for all Figures.

**Table S1.** Comprehensive statistics for Figures 1 and S1.

**Table S2.** Comprehensive statistics for Figure S3.

**Table S3.** Comprehensive statistics for Figure S5.

**Table S4.** Comprehensive statistics for Figure S6.

**Table S5.** Patient cohort information.

**Table S6.** Flow and mass cytometry antibody information.

**Figure S1 (related to Figure 1).**
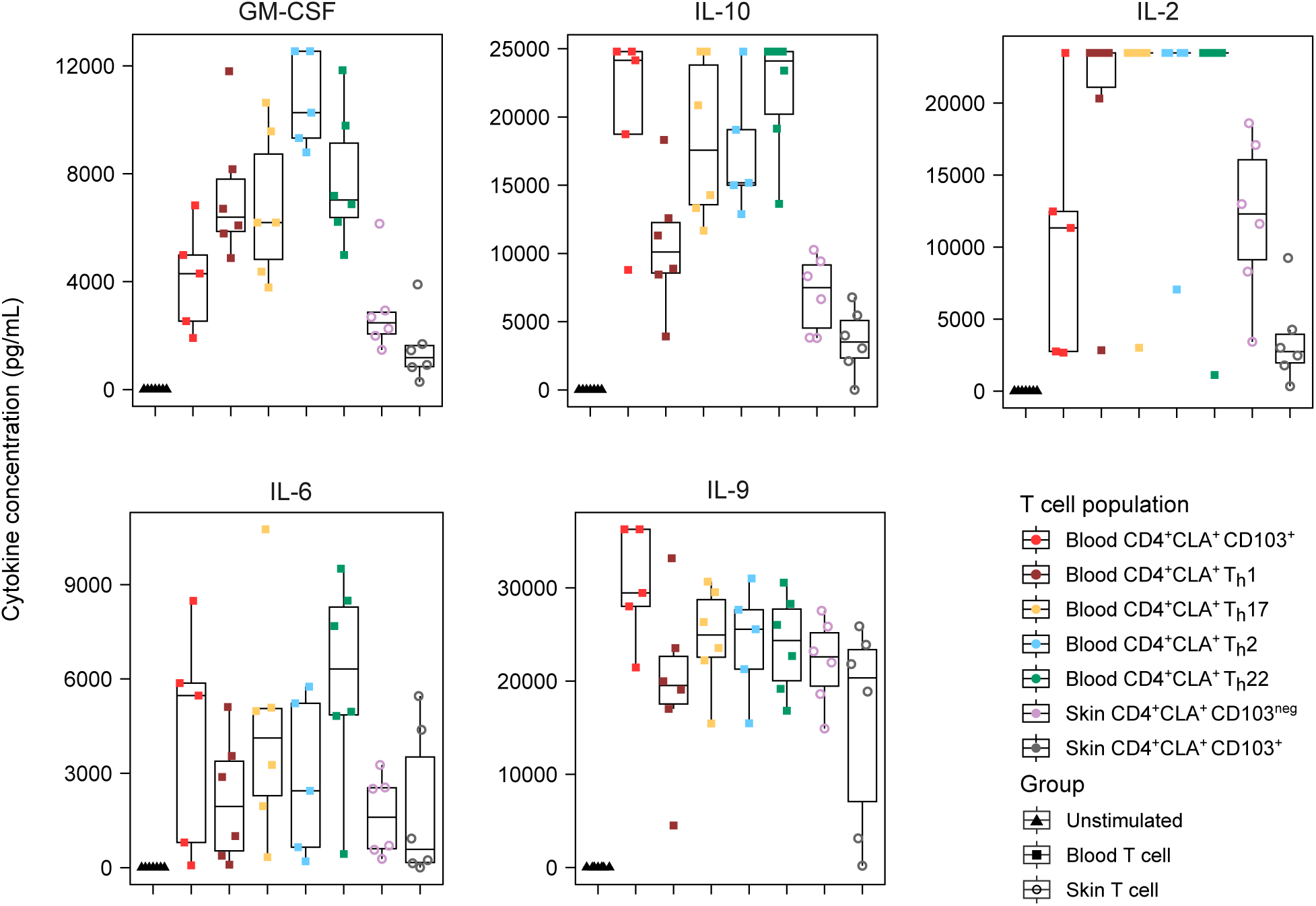
Additional cytokine production capacity by circulating and skin-resident human CD4^+^CLA^+^ T cell populations. Quantitative cytokine bead array measurement of all stimulated T cell supernatants for the 5 indicated analytes (n=5-6 donors per population). Full statistical analysis of cytokine production is provided in **Table S1**.

**Figure S2 (related to Figure 2).**
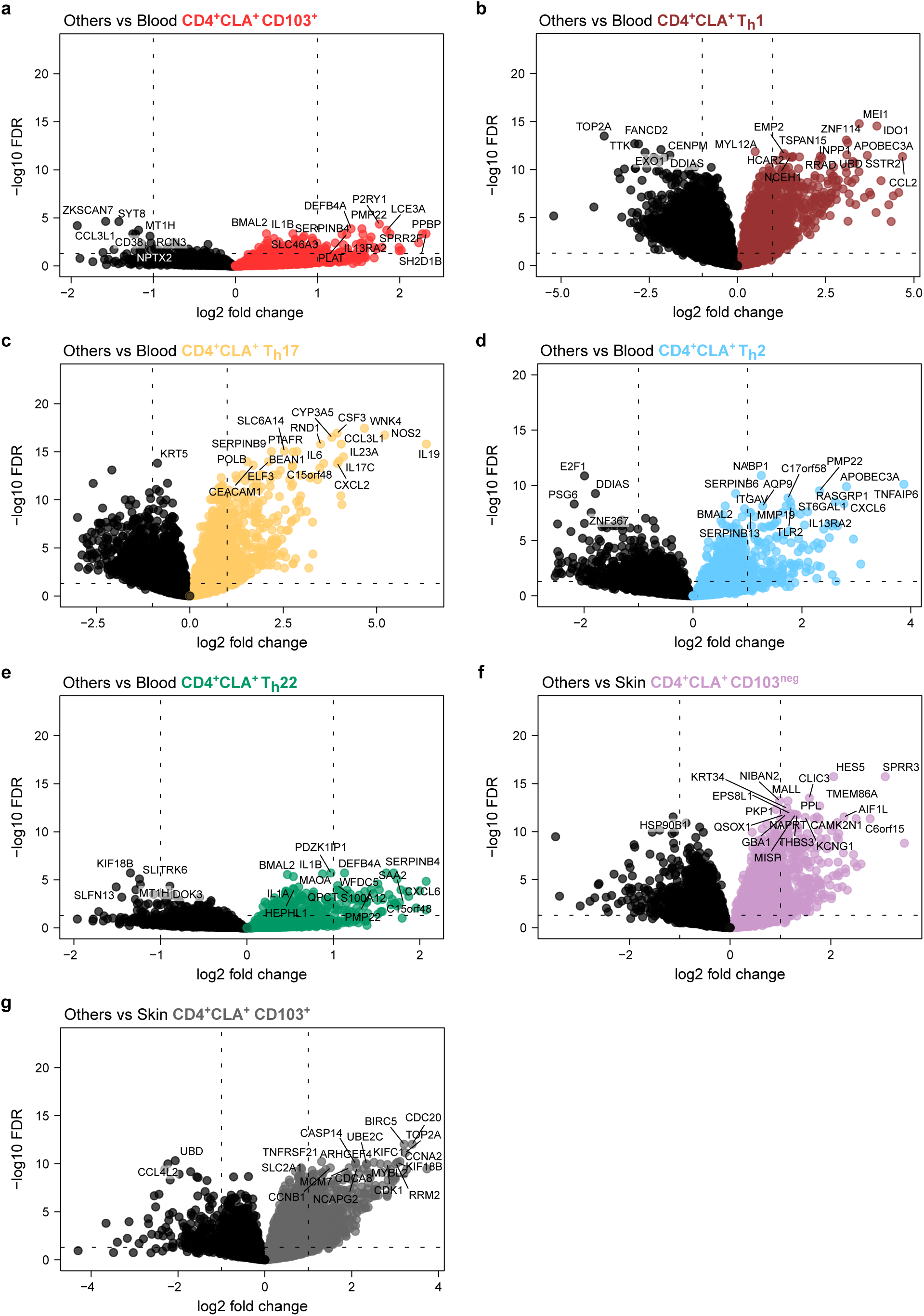
CD4^+^CLA^+^ T cell-induced differential gene expression analysis of epithelial keratinocytes. (**a-g**) Volcano plots showing differential gene expression analysis for KCs stimulated with each indicated CD4^+^CLA^+^ Th population. Cutoffs: log2 fold change >1 and −log10 FDR <0.05. The top 20 genes, ranked by FDR, are labeled on each plot.

**Figure S3 (related to Figure 2).**
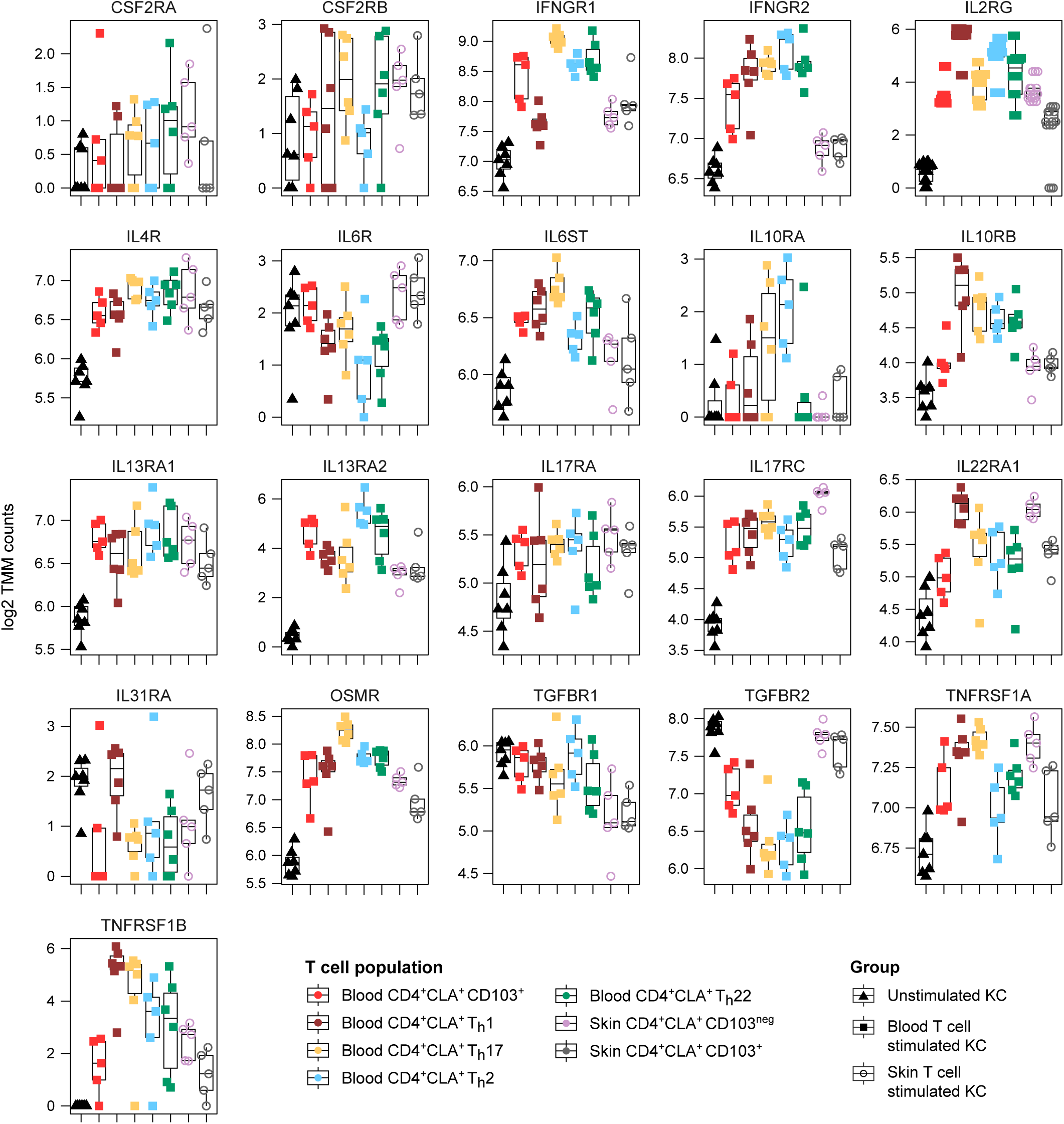
Cytokine receptor gene expression changes induced by CD4^+^CLA^+^ T cells in epithelial keratinocytes. Plots of log2 normalized expression counts using trimmed mean of M values (TMM) for each cytokine receptor gene in KCs stimulated with the indicated T cell populations (n=5-6 donors per population). Full statistical analysis of cytokine receptor gene expression is provided in **Table S2**.

**Figure S4 (related to Figure 3).**
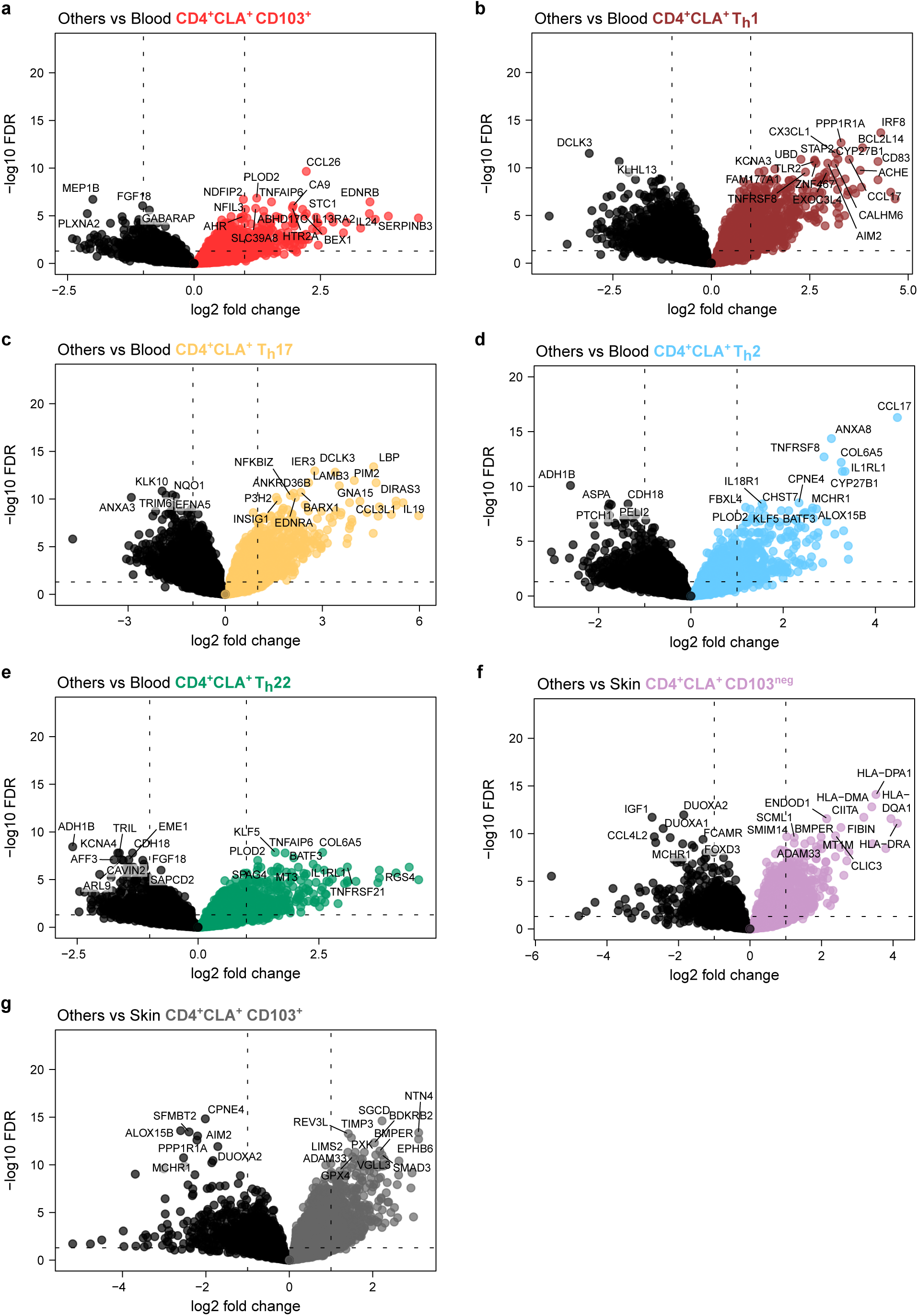
CD4^+^CLA^+^ T cell-induced differential gene expression analysis of dermal fibroblasts. (**a-g**) Volcano plots showing differential gene expression analysis for Fibs stimulated with each indicated CD4^+^CLA^+^ Th population. Cutoffs: log2 fold change >1 and −log10 FDR <0.05. The top 20 genes, ranked by FDR, are labeled on each plot.

**Figure S5 (related to Figure 3).**
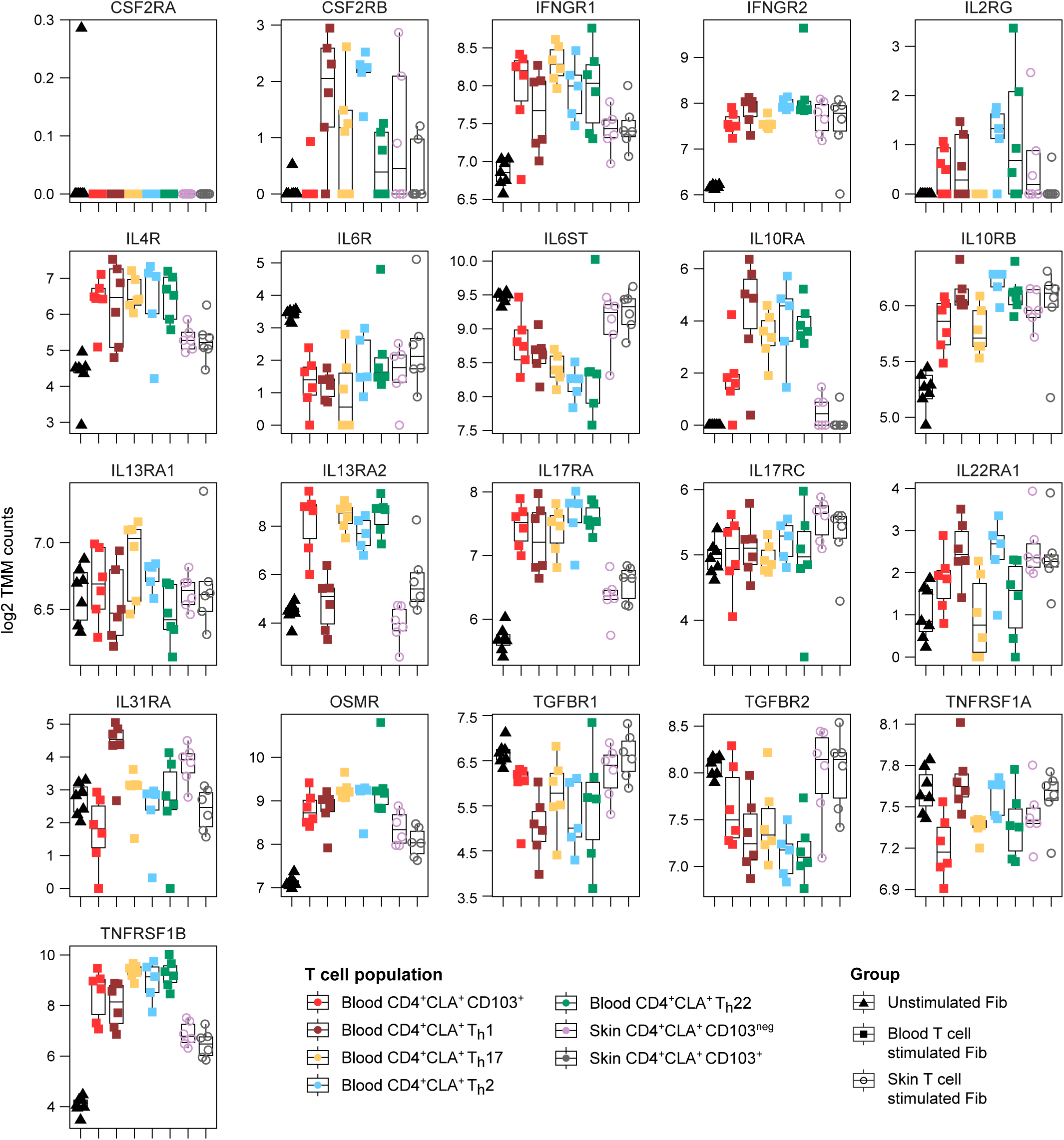
Cytokine receptor gene expression changes induced by CD4^+^CLA^+^ T cells in dermal fibroblasts. Plots of log2 normalized expression counts using trimmed mean of M values (TMM) for each cytokine receptor gene in Fibs stimulated with the indicated T cell populations (n=5-6 donors per population). Full statistical analysis of cytokine receptor gene expression is provided in **Table S3**.

**Figure S6 (related to Figure 4).**
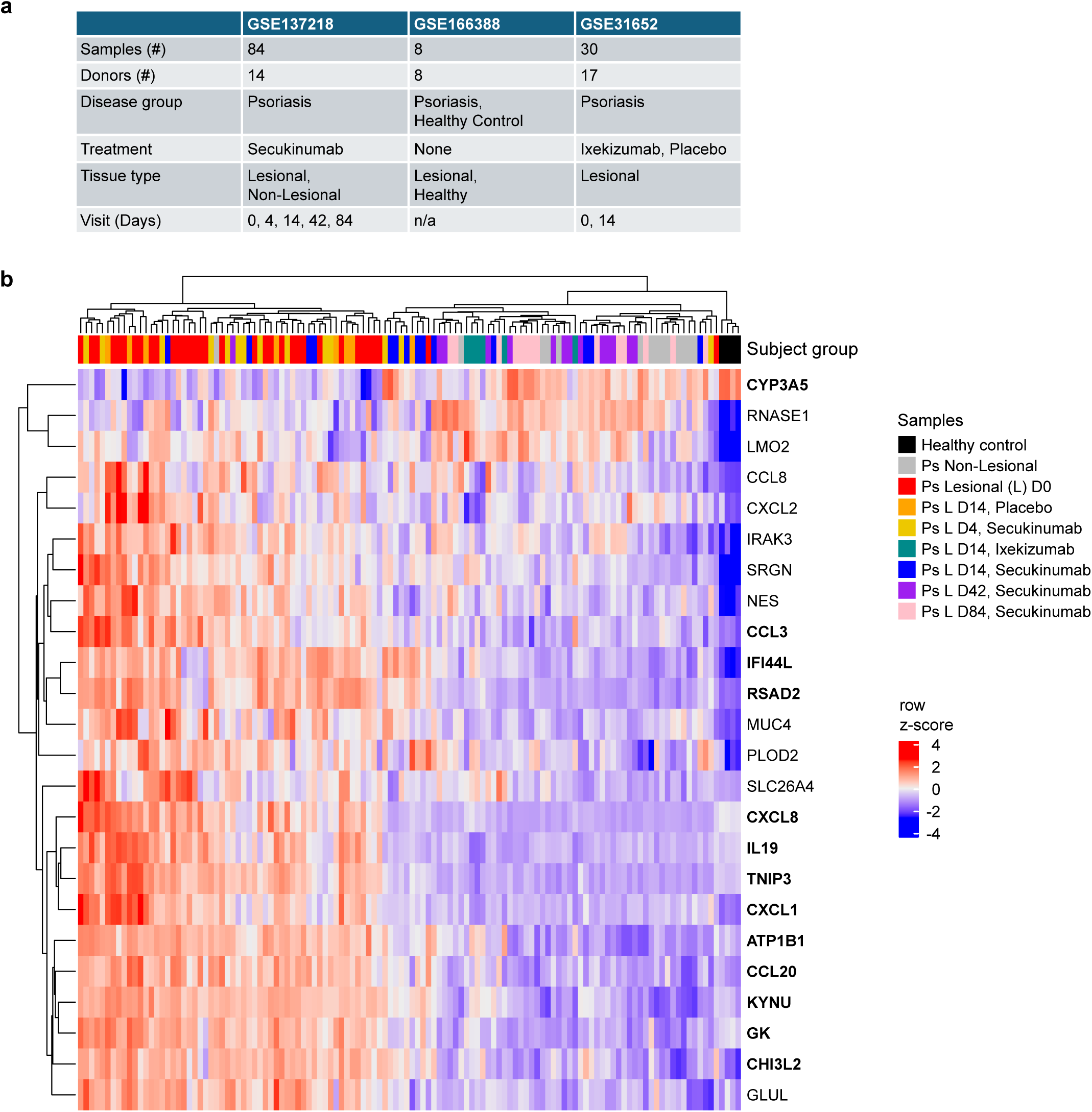
CD4^+^CLA^+^ Th17-dependent genes are altered in psoriasis patients undergoing anti-IL-17A therapy. (**a**) Summary table showing cohort information for each of the three public Ps clinical trial data sets analyzed in Figure 4. (**b**) Heat map showing z-score expression changes of KC genes determined to be Th17-dependent (Figure 2) within publicly available Ps patient data shown in Figure 4 for the indicated groups. Bolded genes are those that are significantly impacted by anti-IL-17A therapy compared with Ps Lesional D0. Full statistical analysis of Th17-dependent gene expression is provided in **Table S4**.

**Figure S7 (related to Figure 5).**
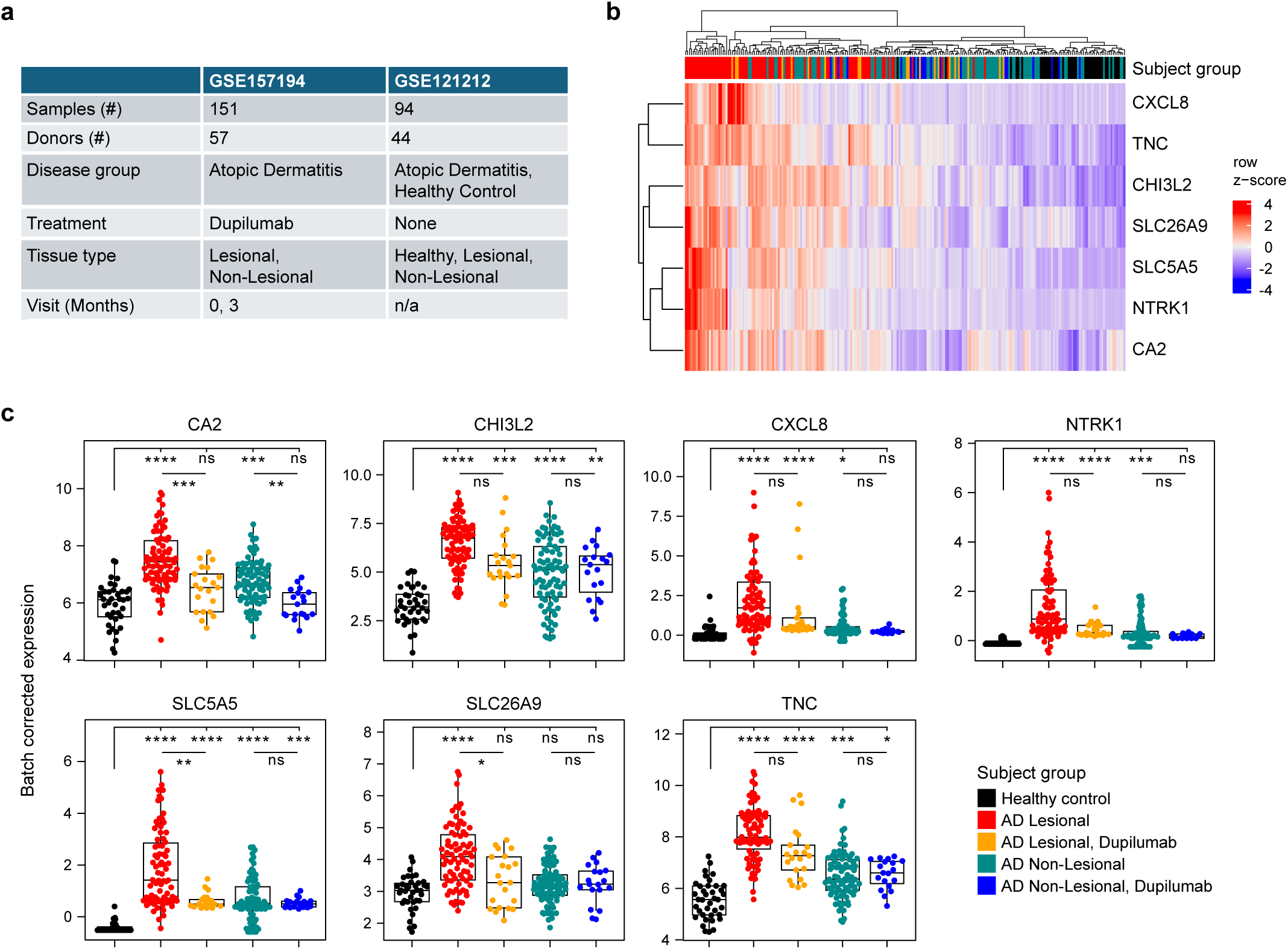
CD4^+^CLA^+^ Th2-dependent genes are altered in atopic dermatitis patients undergoing anti-IL-4Rα therapy. (**a**) Summary table showing cohort information for the two public AD clinical trial data sets analyzed in Figure 5. (**b**) Heat map showing z-score expression changes of KC genes determined to be Th2-dependent (Figure 2) within publicly available AD patient clinical data shown in Figure 5 for the indicated groups. (**c**) Individual gene plots for each Th2-dependent KC gene identified within public AD clinical trial data sets. Error bars indicate mean ± SD; ns = not significant, *p ≤ 0.05, **p ≤ 0.01, ***p ≤ 0.001, and ****p ≤ 0.0001 (Student’s *t*-test).

**Figure S8 (related to Figure 6).**
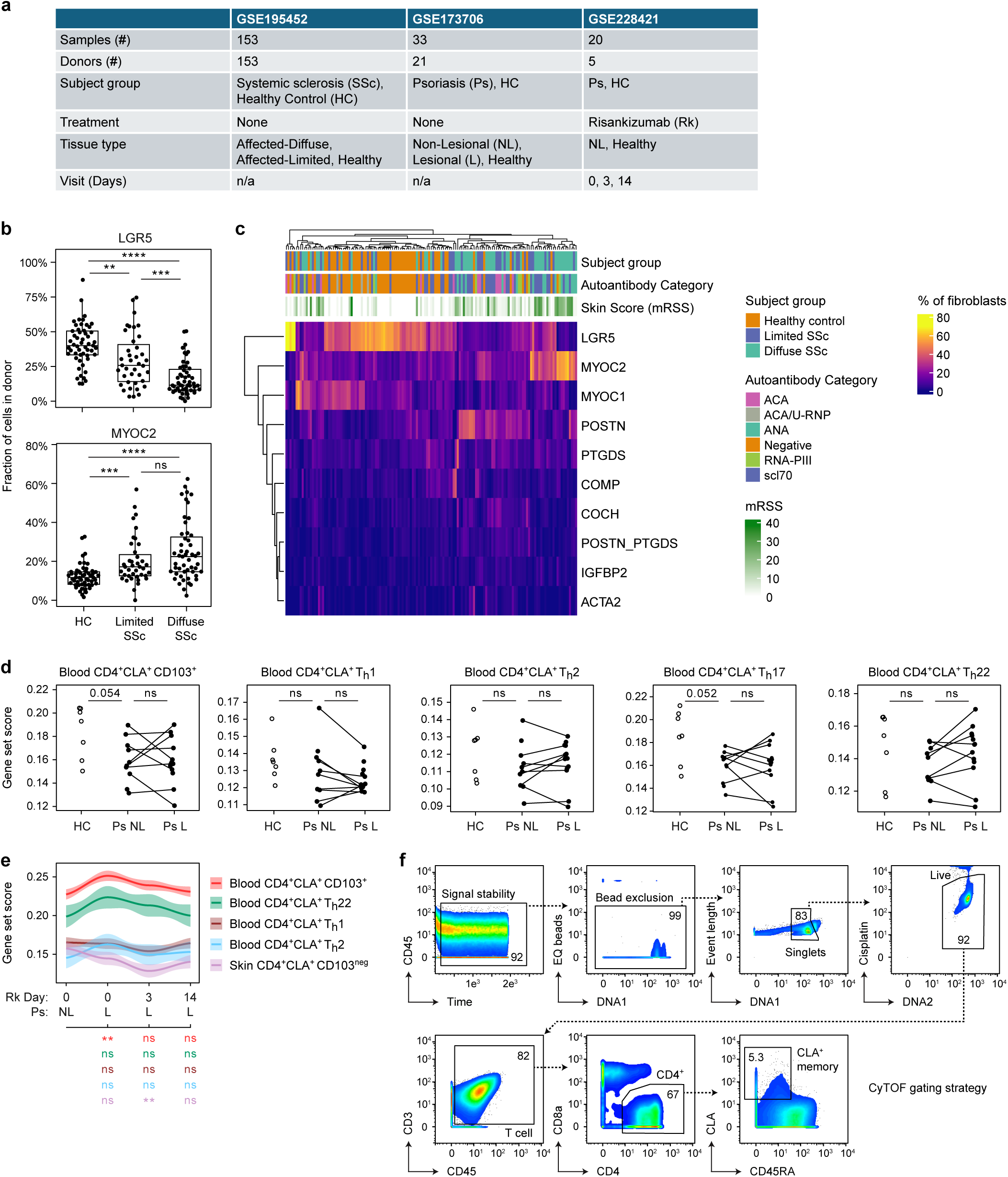
Dermal fibroblast subsets and T cell-dependent fibroblast gene programs are altered between healthy control and inflammatory skin disease subjects. (**a**) Summary table showing cohort information for public data sets analyzed in Figure 6. (**b,c**) Analysis of public scleroderma skin scRNA-seq data [GSE195452]. (b) Plots show the frequency of disease-altered Fib subsets for each subject group. Single cell data are pseudobulked at the subject level. (c) Heat map shows indicated Fib subsets as a percentage of total dermal fibroblasts (*rows*) for each donor (*columns*) including Group, Autoantibody Category, and Skin Score (mRSS). ACA, anti-centromere antibody; U-RNP, U1 small nuclear ribonucleoprotein particle; ANA, anti-nuclear antibody; Neg, auto-antibody negative; RNA-PIII, RNA polymerase III; scl70, scleroderma 70kDa/DNA-topoisomerase-1; mRSS, modified Rodnan skin score. (**d**) Quantification of T cell-dependent Fib gene set enrichment per subject group for public single cell Ps skin data [GSE173706]. (**e**) Indicated T cell-dependent gene set enrichment in single cell skin data of each subject group [GSE228421] undergoing treatment (Tx) with Risankizumab (α-IL-23). (**f**) Mass cytometry identification strategy for CLA^memory^ T cells in donor blood. Error bars indicate mean ± SD; ns = not significant, *p ≤ 0.05, **p ≤ 0.01, ***p ≤ 0.001, and ****p ≤ 0.0001 (Student’s *t*-test).

## Notes

### Competing Interest Statement

The authors have declared no competing interest.

### Summary of Updates

Updated Title; Edits to Abstract, Results, Discussion; New data in Figure 6 & corresponding Figure S8; Updated Materials & Methods; New references; New patient cohort table; New Graphical Abstract

